# 3,500 years of sheeppox virus evolution inferred from archaeological and codicological genomes

**DOI:** 10.64898/2026.02.23.707469

**Authors:** Louis L’Hôte, Luisa Sacristán, Róisín Ferguson, Alex Siekmann, Leland Rogers, Barbara Richter, Herbert Weissenböck, Jenny Lorke, Lara Artemis, Élodie Lévêque, Richard Hark, Patricia Engel, Mary Teresa Josephine Webber, Madison Bennett, Kristine Rose-Beers, Emma Nichols, Marta Munoz Alegre, Max Ramsøe, Sarah Fiddyment, Laura C. Viñas-Caron, Dimitry V. Papin, Ian C. Light-Maka, Felix M. Key, Jonas Albarnaz, Tim Downing, Jonathan Pekar, Philippe Lemey, Daniel Bradley, Jiří Vnouček, Cheryl A. Makarewicz, Joanna Story, Matthew Collins, Matthew D. Teasdale, Annelise Binois-Roman, Sébastien Calvignac-Spencer, Kevin G Daly

## Abstract

Sheeppox virus (SPPV) is a major livestock pathogen causing economic hardship through reduced production and death of vulnerable sheep, with written descriptions of sheeppox-like disease recorded since antiquity. We report 21 novel ancient SPPV genomes spanning the Eurasian steppe Bronze Age (∼1,700 BCE) to the Early Modern period in Western Europe, including multiple genomes obtained from medieval parchment. We estimate that major capripoxvirus lineages diverged ∼11,500-3,700 years ago, overlapping known translocations and bio-cultural developments in sheep. Our dataset supports SPPV diverging first within the lineage leading to goatpox virus and lumpy skin disease virus, and that known gene inactivation events within SPPV and goatpox virus occur in our earliest SPPV genomes. These findings reveal that the food security of Eurasian communities have been threatened by sheeppox for over 3,700 years, and provide new insights to the genomic evolution and potential host adaptation of sheeppox virus.

**Teaser:** Ancient viral genomes from manuscripts and teeth illuminate the evolutionary history of sheeppox virus.

## Introduction

Sheeppox is a highly contagious viral disease that can devastate domestic small ruminant herds, leading to major economic losses for households and communities alike by dramatically reducing milk, wool, and meat production, and often causing death in affected animals (*1*–*3*). Sheeppox-stricken animals typically are visibly ill, the disease presenting with fever, anorexia, ocular discharge, and papular/vesicular lesions on non-woolly areas of the body, in addition to affecting internal organs such as the mouth, lungs, trachea, and digestive and urogenital mucosae (*4, 5*). Morbidity rates can reach 75–90% in immunologically-naïve herds, with mortality rates of around 50% and up to 100% in young animals (*3, 6*). Because of its severity, it has been designated as a notifiable disease by the World Organisation for Animal Health (WOAH). Although eradicated from Europe in the last century, sheeppox remains endemic in Africa, Southwest and Central/Eastern Asia (*6, 7*). It is transmitted primarily through direct contact between infected and healthy animals, but also via aerosols (*8*).

The causative agent of sheeppox is the sheeppox virus (SPPV; *Capripoxvirus sheeppox*), a double-stranded DNA virus belonging to the family *Poxviridae*, the same viral family that includes the variola virus (VARV; *Orthopoxvirus variola*), the etiological agent of smallpox in humans. While smallpox belongs to the genus *Orthopoxvirus*, SPPV is classified within the genus *Capripoxvirus*, together with goatpox virus (GTPV; *Capripoxvirus goatpox*) and lumpy skin disease virus (LSDV; *Capripoxvirus lumpyskinpox*). *Capripoxvirus* genomes are approximately 150kbp in length (*9, 10*), comprising an internal section of well-conserved genes bracketed by ∼2kb terminal inverted repeat regions containing genes associated with host range and virulence (*11*). Capripoxviruses have been assumed to have narrow host tropism, with SPPV infecting sheep, GTPV infecting goats, and LSDV primarily infecting cattle and domestic water buffalo. SPPV and GTPV, which both transmit primarily via aerosols and infected secretions, have occasionally been reported to cause mild infection in goat and sheep, respectively (*2, 12, 13*). In contrast, LSDV transmits mechanically via arthropod vectors and is sporadically detected in wild species such as kudu, giraffes, and Arabian oryx, but has not been reported to affect sheep or goats in natural contexts (*6, 14*). In addition, LSDV has likely been restricted to sub-Saharan Africa until recent decades, whereas SPPV and GTPV have been sampled more widely across north Africa and Eurasia (*2*).

The three *Capripoxvirus* members, SPPV, GTPV and LSDV, are closely related at the genome level, exhibiting a nucleotide similarity of approximately 97% (*15*). However, the phylogenetic relationships within the clade remain unresolved, as different studies have reported conflicting topologies depending on the sequences analyzed and choice of rooting approach; SPPV (*16*), GTPV (*17*) or LSDV (*18, 19*) have each been inferred to be the earliest diverging lineage. Comparative genomic analyses have shown that all genes present in SPPV and GTPV are also found in LSDV. However, a set of LSDV genes (n=8) are inactivated and highly fragmented in both SPPV and GTPV (*15*). A parsimonious explanation of these gene inactivations is that SPPV and GTPV form a clade which is sister to LSDV; that is, a SPPV-GTPV lineage emerged once from a shared ancestor with LSDV (*15, 20*).

Capripoxviruses differ substantially in their earliest occurrence in the historical record, perhaps pointing to a staggered evolutionary emergence. The earliest credible recording of a disease evoking LSDV is dated to 1929 from Zambia (*21*), while the earliest likely documentation of GTPV is from Norway in 1879 (*22*). In contrast, written evidence of SPPV infections appears earlier. Roman sources from the first century CE describe outbreaks in sheep characterized by skin lesions or “pustules”, symptoms consistent with the clinical signs of sheeppox (*23*). During the Middle Ages, references to deadly outbreaks of “pockes” or of “variola” in sheep became increasingly common and are frequently recorded in agricultural treatises and accounting documents from the 13th-16th centuries in France, Spain, and England (*23, 24*). In France, written sources suggest a possible decline in outbreak frequency during the 17th century, followed by a re-emergence in the 18th and 19th centuries, with records of repeated epizootic events and high mortalities. The hiatus appears longer in Great Britain, where mentions of the disease cease by the late 16th century; sheeppox is considered a new and foreign disease when it is reimported with infected sheep in 1847 (*23, 25, 26*). Despite this well-understood recent history of sheeppox in Western Europe, its deeper past is currently unknown.

Here we report the detection of SPPV in 39 Eurasian archaeological and historical specimens, including sheep teeth recovered from archaeological sites in the Eurasian steppe, and parchment and paper specimens from medieval Europe. Our viral detections document SPPV by at least ∼3,800 years ago and are notable in their preponderance within ∼700-1,450 CE, corresponding to the production dates of associated manuscripts. We additionally find SPPV among parchment made from cattle and goat skin, suggesting either cross-species infections or viral contamination from another source. We construct a genome dataset of 22 unique sequences (21 reported here for the first time) of at least 0.1× mean genome coverage (median: 1.5×), including 9 genomes with >5× coverage.

## Results

### Sheeppox virus DNA detected from diverse archaeological and historical remains

To explore the presence of capripoxviruses in ancient livestock through time, we screened 754 samples, both newly reported and previously published (*27*–*39*) from diverse materials spanning over 14,000 years across Eurasia. We identified 41 samples with at least five unique reads (and 3 reads when screening data available were less than 15 million reads) assigned to *Capripoxvirus* using KrakenUniq (*40*) (Tables S1 and S2). In all cases, the best matching source was SPPV, based on comparisons of a *k*-mer based E value between *Capripoxvirus* members (see Methods, Table S1, Figure S1). We then shotgun sequenced samples with higher SPPV DNA representation, and selected a subset of samples with lower SPPV read representation for in-solution RNA bait enrichment of *Capripoxvirus* DNA (see Methods). Merging shotgun sequencing and captured data, we compiled a dataset of 22 unique prehistoric or historic SPPV genomes with more than 0.1× depth (Figure 1, Table S1), ranging from 0.1× to 260.6× and covering at least 10% of the viral genome, including one previously reported (*27*).

**Figure 1:**
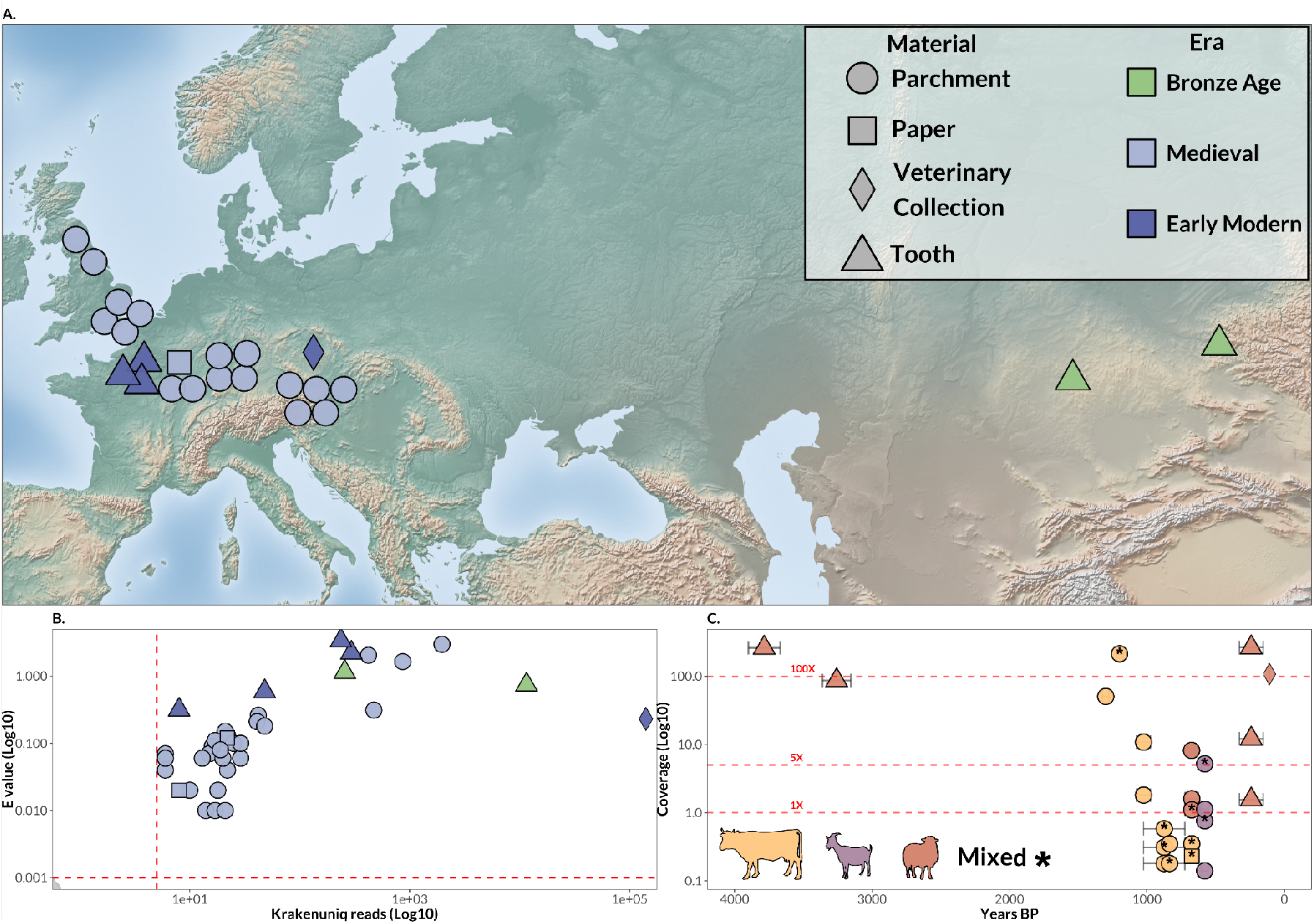
Detection of ancient Sheepox virus genomes in Western Eurasia. A Spatial distribution of all SPPV genomes reaching at least 0.1× of coverage analyzed in this study. Symbols indicate the source type of the SPPV genome, while colors denote archaeological or historical period. B Log10 scale of *k*-mer-based E value relative to log10 scale of the number of reads detected for SPPV by KrakenUniq. Red dashed lines indicate the thresholds used for downstream inclusion (E value ≥ 0.001 and read number ≥ 5). C Genomic coverage (log10 scale) of SPPV-identified samples with ≥ 0.1× mean genome coverage plotted against sample age (years BP). Colors correspond to host species, and shapes correspond to sample material, as in panel A.

Our earliest detections of SPPV occur in Bronze Age pastoralist contexts in the Eurasian steppe. We find two instances of SPPV in sheep teeth from the Late Bronze Age settlement Myrzhik, Kazakhstan (specimen MYRZ180, 1875-1645 BCE, 2σ calibrated range, Table S1) and the Late Bronze Age settlement Rublevo-VI located in the Altai Krai, Russia (RUB002, 1336-1130 BCE, 2σ calibrated range, Table S1). From these, we produced high-quality genomes of 260.6× (MYRZ180, shotgun sequencing) and 86.5× (RUB002, hybrid capture). The settlements from which these SPPV genomes derive represent pastoralist economies centered on intensive sheep husbandry, with flocks managed for both meat and fiber production through livestock management strategies designed to maintain resilient herd structures that buffer against catastrophic loss.

Next, we detected SPPV from numerous parchment and paper-based manuscripts from medieval Northwest and Central Europe – including Britain, the Upper Rhine region, Austria and France (*n* = 17 and 1, for parchment and paper-derived detections respectively, Figure 1 and Table S1). The production dates of the parchment documents span the early 8th century to the 14th century CE, which we assume closely tracks the time of death of the animal whose skin constitutes each parchment page. Our discovery of SPPV occurs in high status codices, including the York Gospels (York, Minster Library, MS Add. 1), a high-status book probably made in Canterbury c. 990–1020 CE, which yielded three SPPV-positive folia, two from the original part of the book and one from a later medieval document appended to it. Other SPPV-positive early manuscripts include the Corpus Glossary (Cambridge, Corpus Christi College, MS 144), made c. 800 CE, again probably in Canterbury, and an annotated copy of the Epistles of St. Paul (Cambridge, Trinity College, MS B.10.5) copied about a century earlier, c. 700 CE. Four SPPV-positive folia were recovered from a portion of the *Speculum historiale*; although the work is a relatively common medieval manuscript, the specific volume analyzed is notable for its historical association with the Vinland Map (*41*–*43*).

Surprisingly, the SPPV-positive manuscript samples derive from membranes of diverse species: we detect SPPV from 10 calfskin folia, 4 from goatskin, and 3 from sheepskin (Tables S1, S3, Figure 1 and Figure S2; see Methods for species assignment based on mitochondrial DNA). The manuscript specimens in which we find SPPV genomes include 8 parchment samples with minor but substantial fractions of animal DNA assigned to additional livestock species (Table S3, Figure S2), and a paper manuscript specimen (“TP01”, Troyes, Médiathèque du Grand Troyes, MS 774). We perform localized sampling on one of these SPPV-positive parchments, TCCL02, and find tentative evidence of greater SPPV yields from the flesh (internal) side compared to the hair (external) side (Table S4). However, these differences were not statistically significant (Wilcoxon rank-sum test, p= 0.250) (Figure S3). In contrast, a fibre sample (TCCL09) yielded a high number of sequencing reads compared to the eraser samples, but most of these reads were associated with the host genome, with only a single read classified as SPPV, indicating poor viral detection despite high sequencing depth.

Our most recent SPPV genomes from archaeological samples originate from a mass mortality assemblage of sheep in the village of Louvres, Val d’Oise, France. The archaeological and palaeopathological assessment of the assemblage, and the genomes themselves, have been detailed elsewhere (*24*). Dated to the 17th–19th century CE, this burial of nine sheep produced four specimens which tested positive for SPPV, in DNA obtained from teeth roots (Table S1). Notably, screening of the corresponding petrous bones did not reveal any SPPV reads, suggesting that SPPV’s occurrence within the respiratory system, oronasal tissue, and within the circulation system (*44*) can lead to long term retention of viral DNA within dental remains, as apparent from the Bronze Age SPPV genomes reported here. Finally, one pathological skin sample from a sheep infected with sheeppox in 1917, preserved in the Veterinary Pathological Museum of Vienna, also tested positive for SPPV.

The authenticity of our recovered SPPV genomes was supported by the fact that pairwise similarity values for all mapped reads were consistently higher for SPPV than for LSDV or GTPV (Table S1). Aligned reads additionally exhibited characteristic ancient DNA misincorporation patterns, with at least 10% of 5′ C→T substitutions at the first base position (Table S1), together with a descending edit-distance profile (r^2^log > 0.60, Table S1, Figure S4), both indicative of genuine ancient SPPV DNA. Additionally, in all but two cases the median aligned read lengths of screened data were below 100 bp (Table S1). Together, these suggest the recovered genomes derive from authentic ancient SPPV viruses.

### Reconstruction of sheeppox virus phylogeny reveals medieval European clade

We then explored the phylogenetic dynamics of SPPV over the last 3,700 years. After constructing a phylogeny of SPPV genomes rooted on GTPV and LSDV, we identified the recovered sequences as SPPV, to the exclusion of GTPV and LSDV (Figure 2A). We additionally identified our oldest genome, MYRZ180 (Myrzhik, Kazakhstan, dated to ∼3,700 BP, Table S1) as basal within the analyzed SPPV sequences. We then placed low-coverage consensus sequences (< 5× coverage) (Figure S5) into the ML topology using pathPhynder (*45*). In addition to our basal MYRZ180 genome, the second SPPV genome recovered from a Bronze Age specimen (RUB002 from Rublevo-VI, Russia) also falls close to the root (Figure 2B) with a relatively short branch length, suggesting it was closely related to the ancestor of all sampled ancient and modern SPPV, which existed just prior to ∼1,200 BCE. As our SPPV dataset shows a clear temporal signal (R^2^ = 0.96; log Bayes factor = 4.34; see Methods and Table S5, Figure S6 and S7; (*46, 47*), we construct a time-calibrated phylogeny (Figure 2C) with BEAST v1.10.5 (*48*) to estimate a mean SPPV clock rate of 9.98 × 10^-6^ (95% HPD: 3.77 × 10^-6^ - 2.31 × 10^-5^). This provides us with an estimate of the coalescence date of our SPPV dataset as ∼4,080 BP (95% HPD: 5,137-3,636 BP).

**Figure 2:**
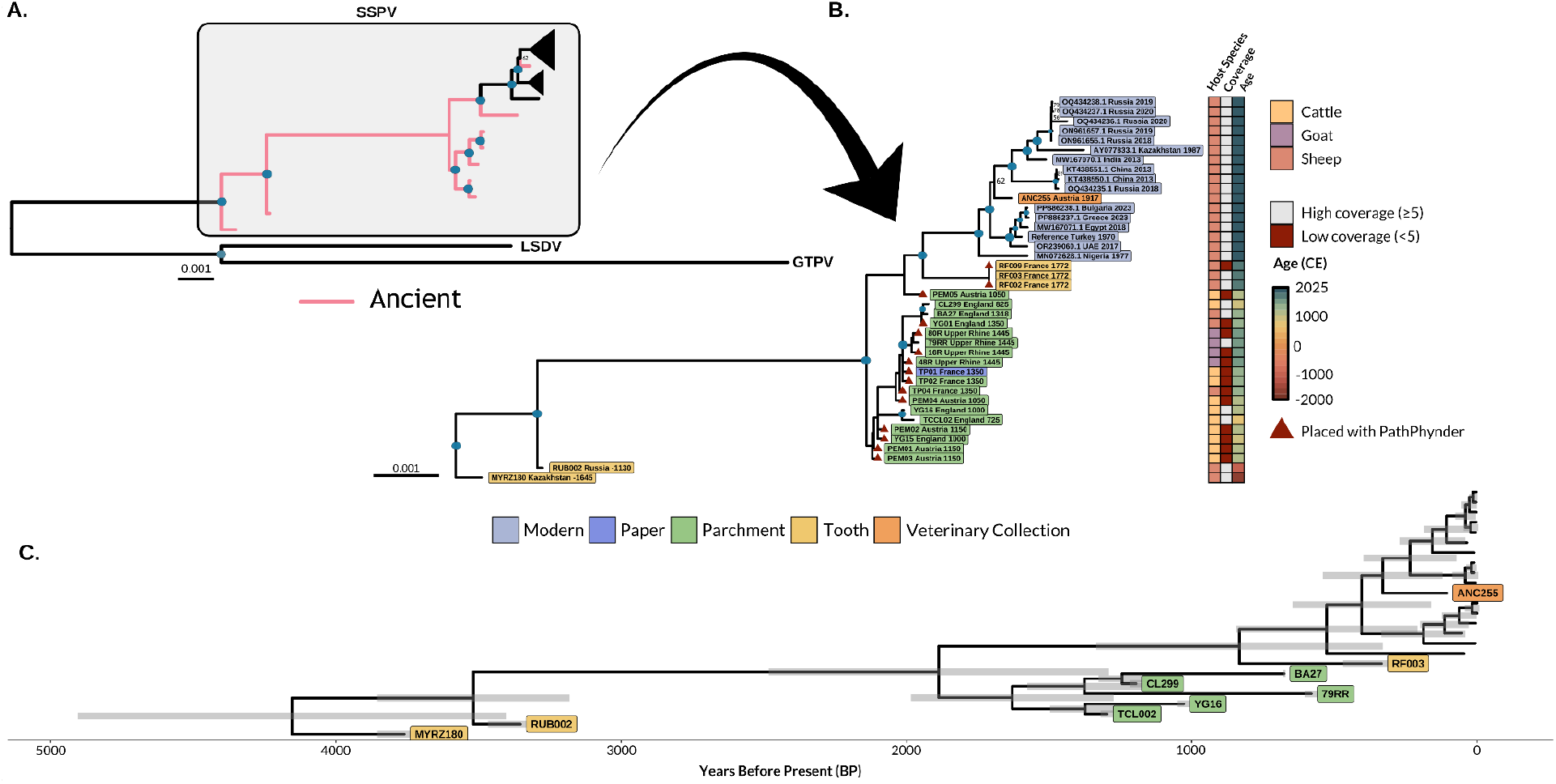
Phylogenetic and dating analysis of ancient and modern sheeppox virus. A. Maximum-likelihood phylogeny of ancient and modern SPPV genomes with ≥5× coverage, rooted using LSDV and GTPV. Nodes with bootstrap support of 100 are indicated by blue circles; other bootstrap values are shown at the corresponding nodes. Clades composed exclusively of modern genomes are collapsed and shown as triangles, and branches corresponding to ancient genomes are highlighted in pink. The grey box delineates the SPPV clade. B. Maximum-likelihood phylogeny of SPPV genomes with placement of low-coverage ancient samples using PathPhynder (*45*). For clarity, only the SPPV portion of the phylogeny is shown. Samples are coloured by material type, and low-coverage genomes (<5×) placed using PathPhynder are indicated by red triangles. Nodes with bootstrap support of 100 are marked with blue circles; other values are shown at the corresponding nodes. Tip labels indicate sampling location and calibrated historical date. Host species, sequencing coverage, and sample age are shown as a heatmap adjacent to the tips. C. Time-calibrated Bayesian phylogeny of the SPPV clade, with samples coloured according to material type.

We observe that all medieval SPPV sequences cluster within, or near, a presumably extinct clade sister to modern SPPV, with potential sublineages detectable among high-coverage genomes (Figure 2B). We estimate the divergence of this European medieval lineage from the ancestor of later SPPV to be ∼1,813 BP (95% HPD: 2,541-1,348 BP; Figure 2C). Lower coverage parchment or paper-derived genomes placed into this scaffold phylogeny do not show a clear relationship between phylogenetic position, provenance, and time. This lack of phylogeographic structure could indicate that multiple SPPV lineages were circulating within Europe during the medieval period; however, we emphasize the limited phylogenetic resolution for lower-coverage sequences. We further hypothesize that our SPPV Beast model could be used to estimate the age of other SPPV-positive parchments of uncertain age; however, we are not able to accurately recover the expected ages of our manuscript-derived sequences, possibly suggesting rate variation among branches of our observed Medieval clade (see Methods and Figure S8). A genome recovered from an 18th century CE sheep mass mortality site in Louvres, France, branches basally to the modern diversity, and is distinct from the medieval parchment clade. Finally, a SPPV genome from 1917 CE, recovered from a sheep skin associated with the Austro-Hungarian Empire, nests within the diversity of modern sequences, all of which coalesce at ∼330 BP (95% HPD: 607-122 BP).

The lack of replacement events between deeply diverged SPPV lineages would similarly suggest gradual, continuous replacement via antigenic drift (*49*). We hypothesized this evolutionary dynamic would be reflected in positive selection within genes producing proteins exposed to the host immune system, such as components of the poxvirus envelope. However, a test for selection in modern (post-1900 CE) SPPV against a background of earlier SPPV failed to find significant signals after correcting for multiple testing (Benjamini-Hochberg FDR) (see Methods and Table S6). It is possible that this reflects limitations in the modern SPPV genomes currently available, which predominantly derive from a small number of outbreaks, and therefore may not reflect longer term evolution within an antigenic drift context.

### Capripoxvirus diversified within the last approximately 5,200 years

We next attempted to resolve the evolutionary history of the *Capripoxvirus* genus, and resolve the sequence of divergence therein. The availability of ancient SPPV sequences provided us with an opportunity to use deep host-specific temporal signals to address the question, albeit at the cost of relying on extrapolations from SPPV lineages, given the lack of ancient GTPV and LSDV genomes. To do this, we applied a Bayesian time-tree approach (*48*) to assess uncertainty in root placement (*50*) within the *Capripoxvirus* clade, using a dataset of 9 ancient SPPV genomes with > 5× coverage, alongside 15 modern SPPV, 9 GTPV, and 32 LSDV genomes (see Figure S9 for a ML phylogeny of this dataset, Table S7). Root posterior probabilities (RPPs) were evaluated under two different clock models: an uncorrelated relaxed clock (*51*), and a shrinkage-based random local clock (*52*) (see Methods). RPPs support a root placement between SPPV and the clade of GTPV and LSDV (RPP=0.44-0.95 across clock models, Table S8, Figures 3B and S10; RPPs of ∼1.0 are obtained when SPPV monophyly is enforced, see Methods). Finally, pairwise similarity values calculated over 500bp sliding windows (100bp steps), excluding ∼4kb potentially affected by recombination and terminal genome regions (see Methods, Figure S11, Table S9), show that in a plurality of windows (467/980, 47.7%), SPPV is more distant from both GTPV and LSDV than GTPV and LSDV are from each other, suggesting that across the genome, SPPV is the most divergent of the capripoxviruses. By comparison, the equivalent tests support SPPV-GTPV and SPPV-LSDV divergence first in 43.2% and 9.0% of windows, respectively. Mean genome-wide pairwise similarly identifies LSDV and GTPV as the most genetically close pair (Figure S12, Table S10).

**Figure 3:**
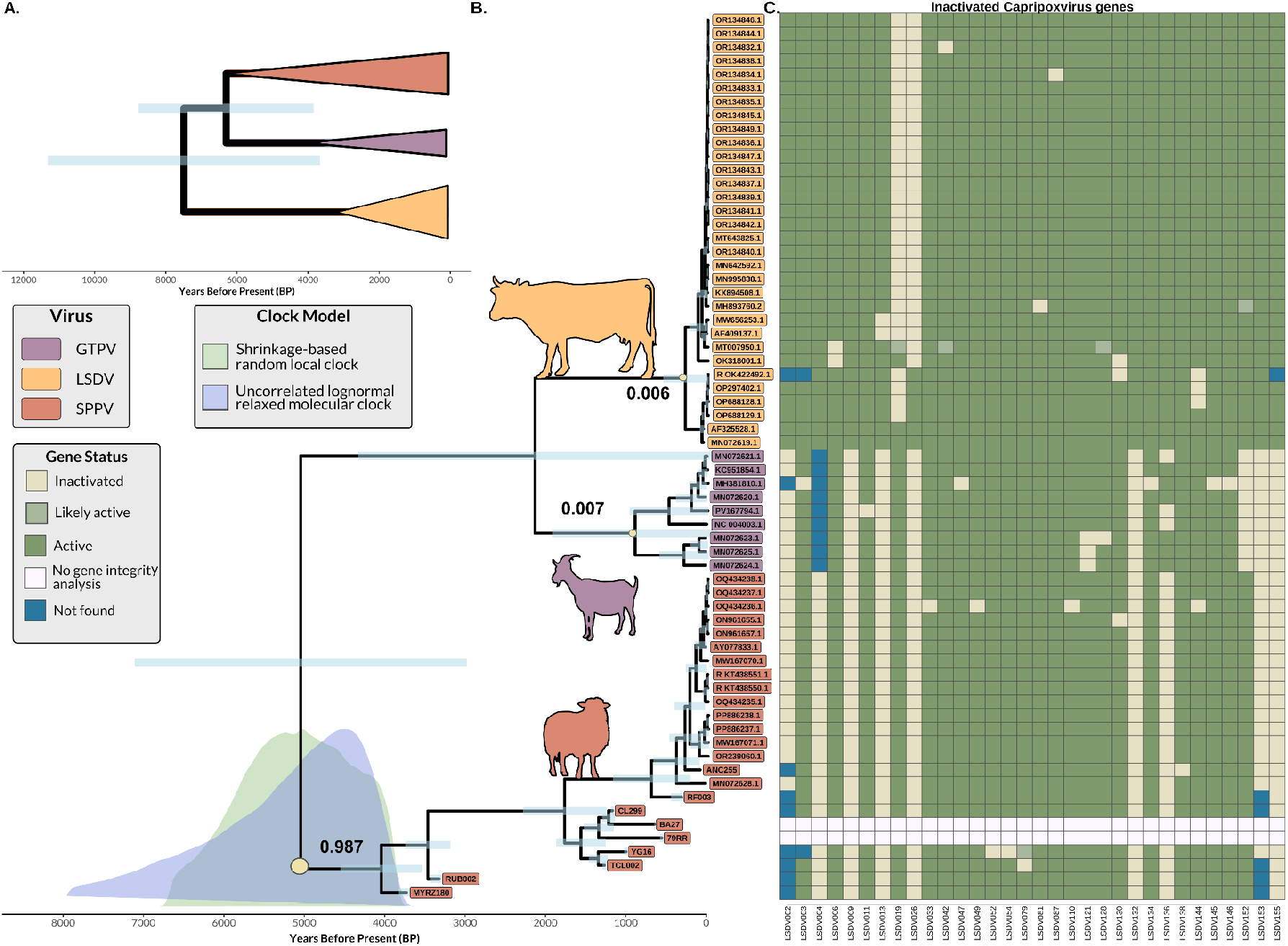
Timing the evolutionary divergence and gene inactivation events of *Capripoxvirus*. A. Time-calibrated Bayesian phylogeny of capripoxviruses with LSDV rooting enforcement. *Capripoxvirus* samples are collapsed at their species TMRCA nodes (and non-collapsed tree is presented in Figure S16). B. Time-calibrated Bayesian phylogeny of capripoxviruses with no rooting enforcement under uncorrelated relaxed molecular-clock model. 95% HPD intervals of the *Capripoxvirus* TMRCA for uncorrelated relaxed molecular-clock model (blue) and the shrinkage-based Random Local Clock model (green) are shown at the root. RPPs for the uncorrelated relaxed molecular-clock model are displayed at major species-level nodes. Tip labels are coloured by viral species, and host silhouettes denote the primary host associated with each lineage. C. Gene integrity matrix of genes which are inactivated at least once in the *Capripoxvirus* lineage, using *de novo* assemblies for the ancient SPPV (two assemblies are excluded due to low quality, see Methods and Table S11). Intact genes are represented in green, likely active in light green, inactivated in yellow, teal if not found in the assembly, and white if the sequence was excluded for the analysis. Columns correspond to viral genes and rows represent genomes included in the time-calibrated Bayesian phylogeny.

Together, these analyses suggest an initial divergence of SPPV within *Capripoxvirus*. However, a ML phylogeny of the core genome rooting on nine other *Chordopoxvirinae* species marginally supports LSDV to diverge first with *Capripoxvirus*, albeit with all other chordopox highly diverged from the Capripoxvirus genus (Figure S13), potentially reflecting long branch attraction (*53*). Additionally, our analyses cannot account for substantial rate variation in the LSDV and GTPV lineages, which could only be detected with the addition of ancient genomes from those clades. Given this inherent limitation, and the observation that SPPV and GPTV share the same pattern of gene inactivation (which could support their sistership), we perform further analyses using two alternative topologies: that either SPPV or LSDV is the outgroup lineage within *Capripoxvirus*.

Our calibration of the *Capripoxvirus* tree provides an estimate of the median sheeppox virus divergence date from other lineages as ∼5,001/5,074 BP allowing SPPV under our two tested clock models (95% HPD: relaxed clock 7,794-3,662 BP; shrinkage-based random local clock 6,572-3,789 BP; Figures 3B, S14 and S15). We infer broader but generally similar divergence estimates when LSDV is enforced as the outgroup sequence (median 7,286 BP; 95% HPD 11,456-3,671 BP, using a relaxed clock, Figures 3A and S16). Divergence times for the LSDV and GTPV are tentatively modeled at ∼2,086/2,199 BP (95% HPD: relaxed clock 4,409-9 BP; shrinkage-based random local clock 4,400-1,548 BP, Figure 3B). Divergence estimates for SPPV and GPTV, when using a LSDV outgroup, are older in comparison (median 6,086 BP; 95% HPD 8,606-3,656 BP, using a relaxed clock, Figures 3A and S16). Regardless of the chosen model, these inferences place the emergence of the sheeppox virus lineage and other capripoxviruses in the millennia following the domestication of their host species in the Holocene.

### Rapid gene inactivation within sheeppox virus and parallel inactivation in goatpox virus

Gene loss or inactivation is a recurring genomic pattern associated with the evolution of host specificity in poxviruses (*54*) and more broadly in human-affecting pathogens (*55*). Analysis of ancient VARV genomes has revealed a pattern of progressive gene inactivation over time between ancient and modern VARV strains (*56*). We sought to test whether a similar pattern occurred in ancient SPPV compared with modern viral genomes, and the pattern of gene inactivation more broadly within *Capripoxvirus*. Comparative genome analysis of *Capripoxvirus* has revealed that it consists of 156 genes which are functional in LSDV; 8 of these genes are inactivated both in SPPV and GTPV (*15, 20*) (Table S11), while one gene (LSDV136) is inactivated in SPPV and still active in some GTPV clades.

We then analyzed *de-novo* reconstructed genomes (passing our defined thresholds of assembly quality, Table S12) (MYRZ180, RUB002, TCCL02, CL299, YG16, RF003 and ANC255), all representing distinct time eras, and modern *Capripoxviru*s genomes to time gene inactivation events within their evolutionary history. By annotating and analyzing the integrity of homologous genes between the three *Capripoxvirus* members, we observed that the 9 genes inactivated in modern SPPV are all inactivated in analyzed ancient SPPV genomes (Figure 3C, Table S11), including our oldest genome MYRZ180 (∼3,700 BP). This constrains the timing of these inactivation events to, at the latest, the divergence time of MYRZ180 from other sheeppox viral lineages ∼4,004 BP (95% HPD: 5,107-3,662 BP, Figure 2C). The rapid inactivation of these nine genes 1,600 years after our estimate of the divergence of the SPPV lineage (the median coalescence estimate of *Capripoxvirus* under our relaxed clock, “SPPV outgroup” model) suggests that these genes were central to the adaptation of SPPV to their sheep host. However, unlike VARV and other human-infecting pathogens (*55, 56*), SPPV does not show further gene inactivation through time (Figure S17, Table S11) and seems to have a relatively constant number of active genes since the Bronze Age. If these gene inactivation events underlie (or partially underlie) adaptation to their sheep host, these results suggest the initial adaptation process was already complete by ∼4,000 BP, approximately when our oldest genome coalesces with other SPPV.

We then assessed the sharing of frameshift and nonsense mutations in these inactivated genes ‐ potentially responsible for inactivation ‐ across SPPV and GTPV. Comparison of 354 disruptive mutations found in our datasets falling within the 9 inactivated genes revealed that only a single such mutation is shared by all SPPV and GTPV genomes. This frameshift mutation, located at position 142 of the LSDV026 protein alignment, is likely an alignment artefact, as it was also detected in all LSDV genomes in which the gene is otherwise intact (see Methods, Figure S18, Table S13). The interpretation of this finding is dependent on the identity of the outgroup sequence, with an LSDV outgroup suggesting it is no longer possible to identify the original, shared inactivating mutation between both lineages (presumably because it has been obscured by further decay after inactivation). Alternatively, if SPPV diverged first, the absence of any shared disruptive mutations between SPPV and GTPV would suggest these nine gene inactivation events occurred independently in the two lineages, rather than being inherited from a common ancestor.

## Discussion

Our results extend the molecular evidence for sheeppox by ∼3,400 years (*27*), provide the first genetic detection of SPPV in prehistory, and to our knowledge represent the current oldest genomes of *Poxviridae* recovered. Given the inferred early pseudogenization (Figure 3) and reports of a sheeppox-like disease (“the pustule”) by a Roman author, Columella, writing in the first century CE (*57*), it is plausible that sheeppox was a similarly serious disease at this time. The presence of SPPV in Central Kazakhstan (Myrzhik, specimen MYRZ180, dated 1875-1645 cal BCE) and the Altai Krai region of Russia (Rublevo-VI, specimen RUB002, dated 1336-1130 BCE) indicates that by the Bronze Age period, domestic sheep herds of the Eurasian steppe region were afflicted by a pathogen which today is highly infectious and virulent. *Yersinia pestis*, the bacterial agent of the plague, has also been detected in sheep from this region and period (*34*), suggesting that herding societies in the Steppe may have contended with a diverse burden of zoonotic and livestock-specific diseases.

Sheeppox likely posed a significant threat to food security, household subsistence, and opportunities for surplus production in the ancient steppe for both mobile pastoralists and more sedentary agro-pastoralists alike. Dramatic declines in milk yield, wool and hide quality - the latter compromised by lesions caused by SPPV, and overall animal survivorship would have sharply curtailed the availability of meat, fat, and dairy, impaired production of felts and woven textiles used for shelter and insulation, and reduced supply of skins essential for container and tool manufacture. Moreover, sheeppox- induced mortality would have undermined pastoralist planning of herd security, critical for sustaining flocks in harsh steppe environments, by unpredictably disrupting the scheduled culling of specific age and sex cohorts used to manage herd structure and optimize productivity over the long term. Large settlements in the steppe were key nodes in the exchange of ceramics and metals (*58, 59*). The presence of sheeppox in steppe settlements engaged in intensive metallurgical production hints at how emplaced herds aggregated to provision resident craft specialists may have functioned as enzootic reservoirs of infection. The inflow of ores metal-producing settlements and outflow of finished metal goods was reasonably mediated by mobile pastoralist households undertaking seasonal transhumant movements, which may have facilitated transmission of sheeppox through flock-to-flock contact along exchange and mobility corridors.

Populations in the Northern Caucasian steppe region consumed sheep dairy from the 4th-1st millennium BCE (*60, 61*). Additionally, the East European steppe region provides some of the earliest evidence of preserved sheep wool ∼2,400-2,000 BCE, before becoming more widespread from 2,000 BCE (*62, 63*). Sheep populations from the steppe region were likely translocated to Europe by the mid-2nd millennium BCE, contributing ∼50% of ancestry in later sheep (*25*). In the case that SPPV emerged outside of Europe, this large-scale animal movement event may have had considerable epidemiological consequences to the sheep populations present in Europe at the time. Investigating the presence of SPPV in Europe among ovine remains from before and after the Bronze Age period would help clarify whether the virus was already circulating within Europe.

We additionally discover SPPV within medieval parchment and paper artifacts across Central and Northwestern Europe (Figure 1). To our knowledge, this is the first time pathogen genomes have been recovered from such historical materials, and suggests that other pathogens could be similarly preserved in codicological specimens. The occurrence of SPPV DNA in parchment is likely a product of the high viral titre in skin tissue during active sheeppox cases (*44*). The parchment folios in which we recover SPPV DNA do not show visible lesions (Table S1). This may not be surprising: sheeppox dermatological lesions are often confined to non-wool parts of the body (*4*), and the production of parchment involved the removal of the flesh itself through scraping (*64, 65*). However, there are indications of pox-like lesions on the membrane of the early 8th century CE Codex Amiatinus (Florence, Biblioteca Medicea Laurenziana MS Amiatino 1), made of a combination of goat and sheep skin parchment (*66*). It is also possible that, in the past, sheep skinners actively avoided body regions bearing such lesions when producing hides that were later used for ecclesiastical manuscripts and legal documents, or that affected elements were rejected by scribes. Indeed, an assessment of SPPV-positive sheep from 17th–19th century CE France indicated that some of those animals were flayed despite being affected by sheeppox (*27*), suggesting that in at least some cases, infection would not dissuade the harvesting of an infected animal’s hide.

The detection of SPPV in parchment derived from tissues of species other than sheep, as determined through a competitive mitochondrial DNA analysis (Table S3, Figure S1), is surprising. While SPPV has been reported in a small number of outbreaks in goats (*67*–*69*), to our knowledge there are no reported natural cases of SPPV infections in cattle (*70*). Additionally, parchment was typically produced from calves 6-8 weeks of age (*71, 72*); if cattle had themselves been exposed to SPPV during their lifetimes and developed a humoral response, their calves should be protected via colostral immunity (*73*). The possible recovery of greater SPPV reads from the flesh side rather than hair side during localized sampling (Table S4, Figure S3) of one specific parchment leaf, TCCL02, may support a case of in infection for this cattle skin, which lacks any signal of sheep mtDNA (see below and Table S3).

Alternatively, SPPV DNA or virions may have been transferred to calf and goat folia during the parchment production process. This typically involved soaking flayed skins in a pit filled with a water- lime bath (*64*). Contamination within such pits may have been facilitated by the persistence of the SPPV virion outside the body on the scale of months, particularly from shed scabs (*2*). Another explanation could be the topical contamination of SPPV between parchment in the same manuscript, where adjacent pages were derived from infected sheep and non-infected calves. A final possibility is the use of animal-derived (i.e. sheep) products in the production, repair, or modification of a non-sheep parchment. Indeed, 6 of the 8 parchments with mixed signals have sheep mitochondria-aligned reads as the second-most commonly represented, with another giving a predominantly sheep signal (Table S3). Additionally, our detection of a SPPV genome within a paper document (“TP01”) may also be explained by its production process. As paper in Late Medieval and Early Modern Europe was often sized with animal-derived gelatin to improve its durability and writability, the animal and viral genetic signals may derive from introduced gelatin (*74*–*76*). Indeed, the glue used to size paper was often produced from livestock leather scraps or parchment trimmings (*77*).

Our estimation of the *Capripoxvirus* divergence time places this event conservatively ∼11,500-3,700 years ago (the widest range across our four tested models); rather than evolving with their host species, *Capripoxvirus* members likely diversified in the millennia during or following the onset of domestication of their host livestock (*78*). Ancient genomes from other pathogen lineages, such as measles virus/rinderpest virus (*79*), *Salmonella enterica* Paratyphi C (*80*), *Brucella melitensis* (*33)*, and *Borrelia recurrentis* (*81)*, either diversified or adapted to human or livestock hosts within the last 10,000 years. Mixed livestock farming and the use of animal products may have therefore catalyzed the emergence of pathogen lineages adapted to specific hosts, both human and animal, and could be considered a consequence of domestication. Notably, our SPPV divergence estimates peak roughly 5,200-4,200 years BP, a period overlapping with the inferred major translocation of sheep from the Eurasian steppe to Europe (*28*) and the earliest preserved remains of wool (*63*). The demographic processes underlying either this migratory event or the emergence of woolly sheep could have facilitated the initial transmission and adaptation of SPPV within its ovine host.

While our calibrated analyses unambiguously identify SPPV as the most basal capripoxvirus (Figure 3B), they are inherently limited by the fact that the temporal signal is mostly derived from the ancient SPPV genomes (see Methods, LSDV R^2^: 0.16 and GTPV clades 2.1,2.2 and 2.3 R^2^: 0.06,0.0 and 0.003, Figure S19). Therefore, we cannot exclude that the lineages leading to contemporaneous GTPV and LSDV have evolved at a different rate. If it was the case, the root may be positioned on another branch than the one our data point to today i.e. SPPV. The only way to investigate this hypothesis is to obtain ancient genomes from either LSDV or GTPV. If SPPV is in fact, as suggested here, the outgroup, the tree topology is notably incongruent with both host species relationships, where sheep and goat share a more recent common ancestor (*82*), and with the pattern of gene inactivation across *Capripoxvirus*, where eight genes activated in LSDV are non-functional in SPPV and GTPV (Figure 3C). This evolutionary relationship would imply independent and parallel inactivation of these eight genes in viral lineages affecting sheep and goat. In such a scenario, repeated emergence from a cattle-infecting strain may have caused parallel inactivation within the SPPV and GTPV lineages due to analogous pressures to survive the ovine and caprine immune systems. The recovery of ancient genomes of either GTPV or LSDV may definitively resolve the question of the root of the *Capripoxvirus* phylogeny and, consequentially, the succession of events that led to these viruses’ diversity and distribution in the three bovid species.

In conclusion, our findings provide the first molecular understanding of sheeppox virus in pre-Modern times. Although present in the Bronze Age, we find a particularly high number of SPPV genomes in the medieval period, associated with animal skin-based documents, implying that infection of sheep was a frequent challenge to European societies of this time. Our current data clearly supports an initial divergence of SPPV within *Capripoxvirus*; temporal genome series from other capripoxviruses may cause this topology to be revised. We demonstrate how genomes derived over millennia can chart the emergence and ongoing adaptation of pathogens to their hosts, inferring gene inactivation events which occurred by ∼4,000 years BP. Our study is limited in the geographic regions assessed, the number of dental specimens studied, and by the *Capripoxvirus* members thus recovered. Ancient, historic, or Early Modern genomes from GTPV and LSDV may complement the SPPV genomes reported here - potentially from animal skin-based documentation - and allow for a more complete reconstruction of the evolution of this important genus of livestock pathogens.

## Materials and Methods

### Sample provenance

#### Rublevo-VI

Rublevo-VI was a substantial Late Bronze Age settlement center and metallurgical production site occupied during the mid-late second millennium BCE, situated in the forest-steppes of the Altai Krai (Russia) (*83, 84*). Exceeding 10 hectares in its extent, Rublevo-VI was intensively occupied, reflected in a dense cluster of at least fifteen structures and an expansive midden pit, covering roughly 400 m^2^, purposefully cut into a sand dune adjacent to the structures. The midden, characterized by repeated ash deposition, reddish-brown colored layers and lenses of sooty, sandy loam suggestive of in situ burning events, contained high densities of fragmented ceramics associated with the Sargary-Alekseyevka ceramic tradition, as well as abundant faunal skeletal remains.

Rublevo-VI was also a locus of metallurgical activity evidenced by ore, slag, stone casting molds, and finished metal products as well as metal fragments recovered from the site (*85*). The use of reusable molds further suggests a degree of standardized, possible large-scale production, implying that metal items were likely produced for exchange beyond the settlement itself. Although the extent of agricultural activity at the settlement remains unclear, livestock husbandry focused on cattle (36% of faunal remains) and sheep herding (28%) with occasional use of goats, but intensive use of horses (43%), and occasional hunting of elk (*86*). Young adult sheep were slaughtered for their meat once they had reached peak body weight, while older sheep were retained until around six years of age, likely as part of a herd security strategy aimed at reducing risk to the flock. Altogether, Rublevo-VI appears to have supported a sizable community, serving as a regional economic and cultural center through which ore, metals, ceramics (*83*), and animals flowed.

While previous liquid scintillation counting radiocarbon dating of charred animal bone, sooty loam layers, and buried soils placed occupation of Rublevo-VI during the 12−9^th^ centuries BCE (*87*), the new radiocarbon determination measured from a sheep molar presented in this study suggests an older occupation dating to the mid-second millennium BCE (CIRAM-8056, 3021 ± 30 BP; 1,392-1,130 BCE).

#### Myrzhik

Myrzhik is a Late Bronze Age settlement and major metallurgical production site situated on the Atasu River in the open steppes of Central Kazakhstan. Ceramic chrono-typology places occupation of the site during the 14^th^–8^th^ centuries BCE, but the sheep molar analyzed as part of this study (OxA-45742, 3340 ± 19 BP; 1,875–1,645 BCE) suggests significantly earlier occupation. Although relatively small in size (∼4 h), Myrzhik was densely occupied, with nearly forty semi-subterranean post-and-frame residential and workshop structures visible on the unexcavated surface as circular, rectangular, and double-lobed shaped depressions (*88*). The settlement exhibits two distinct occupation horizons, the earlier distinguished by large semi-subterranean rectangular structures (∼6 x 10m) partially lined with large stone slabs vertically set in an underlying ashy substrate, and featured in their interior a hearth and numerous pits containing ashy fill yielding Alakul-type ceramic fragments, faunal remains, discarded copper ore, as well as items including pestles and bronze awls. The later horizon is characterized by roughly oval semi-subterranean dwellings associated with Sargara-type ceramics. These structures were similarly large, with postholes positioned along the walls and in the center of the dwelling, supporting building superstructure and roof (*88*).

Mryzhik was also a major metallurgical center, supporting multiple workshop areas containing substantial, well-constructed pit furnaces, more industrial-scale ore smelting furnaces including a three-pit configuration encircled by ring-shaped channels extending vertically downward to regular airflow (*89*), and other pit furnaces associated with complex horizontal and vertical channel systems (*88*). The abundance of heavy pestles, grinding stones, fired stones, oxidized copper ore, slag, and copper pieces further attests to intensive smelting, while fragments of bronze objects and small bronze tools and adornments such as awls, arrowheads, bracelets indicate at least some foundry work also took place. Pastoral production was also important at Myrzhik, focused on intensive husbandry of caprines (70% of the total faunal assemblage, NISP = 5503), mostly sheep, further supplemented by cattle herding (30%) (*90*). Sheep were managed for their meat and wool, but notably not for dairy, indicated by intensive kill-off of prime aged young adults and fibre-producing older animals (*91*). Horses were very rarely used.

Myrzhik was excavated from ∼1978 to 1984 by the Central Kazakhstan Archaeological Expedition, directed successively by M.K. Kadyrbayev, Zh. Kurmankulov and S.M. Akhinzhanov, and finally by E.F. Kuznetsova during the 1984 season, when the sheep specimen analyzed as part of this study was recovered; more recent excavations were undertaken in 2016-2017 (*88*). The faunal assemblage recovered from Myrzhik is archived at the Margulan Institute of Archaeology Committee of Science Ministry of Science and Higher Education of the Republic of Kazakhstan.

#### Speculum Historiale

The *Speculum Historiale* (“Mirror of History”) is one section of the thirteenth-century encyclopedic *Speculum Maius* compiled by Vincent of Beauvais. Yale University holds a fifteenth-century manuscript comprising books 21–24 of the *Speculum Historiale*, which was formerly bound together with the Vinland Map and the *Tartar Relation*. This manuscript is generally understood to have been commissioned in connection with the Council of Basel (1431–1449), and radiocarbon dating places its parchment in the early to mid-fifteenth century, consistent with paleographic and codicological evidence.

The bound *Speculum Historiale* and *Tartar Relation* volume was removed from the cathedral library of La Seo in Zaragoza, Spain, in the post–World War II period; ownership stamps at the bottoms of several folios were excised, and the losses were repaired with replacement parchment, including folio 79RR, which tested positive for sheeppox virus in the present study. The Vinland Map was drawn on parchment that was originally part of the *Speculum Historiale* manuscript, likely a flyleaf from the front of the volume, as indicated by the presence of matching wormholes in the Vinland Map and the first pages of text in the *Speculum*.

When the Vinland Map was introduced to modern scholarship in the mid-twentieth century, it generated considerable interest because its depiction of “Vinland” west of Greenland appeared to imply Norse knowledge of North America predating Columbus’s 1492 voyage. The physical association of the map with the *Speculum Historiale* manuscript was therefore cited as evidence for a medieval context. Subsequent multidisciplinary analyses have demonstrated, however, that the ink used to draw the map is of modern origin, establishing the map itself as a modern fabrication.

#### Clairvaux manuscripts

Clairvaux is a Cistercian abbey located in Champagne (present-day Aube, France). Founded in 1115 by Bernard of Clairvaux, it became one of the most influential monasteries in medieval Europe. Clairvaux was a major spiritual, intellectual, and institutional centre of the Cistercian Order, founding dozens of daughter houses across Europe. Its importance rested not only on Bernard’s authority, but also on its scriptorium and library, which grew into one of the largest and best-organised monastic libraries of the Middle Ages.

- MS 774 was copied in Paris between 1450 and 1455 for the abbot of Clairvaux (Pierre de Virey) and belonged to him. Title: *DURANDELLUS, Evidentiae contra Durandum*; *JOHANNES NEAPOLITANUS diac*. Pierre de Virey was the Cistercian abbot of Clairvaux from 1471 to 1496. He is best known for his role in the administrative and intellectual reorganisation of the abbey, particularly its library. Soon after his appointment, he commissioned a thorough inventory of the abbey’s books, resulting in the 1472 catalogue compiled by Jean de Voivre.
- MS 2299 is the original library catalogue, commissioned by the same abbot and written by his librarian Jean de Voivre in 1472. Title: *Inventoire et declaracion des volumes et livres de l’eglise et abbaye de Clervaulx,[*…*] fait au moys de may l’an mil CCCC LXXII, par nous frere Pierre de Virée, nouvel abbé dudit lieu, [*…] (Ms 2299). Pierre de Virey became abbot of Clairvaux in 1471, at a time when the abbey’s library was being reorganised. He soon ordered a formal inventory of the collection. Compiled in 1472 by Jean de Voivre, who was probably acting as librarian or prior at the time, this catalogue offers an excellent insight into the library at the start of Pierre de Virey’s abbacy. It catalogues an exceptionally large and well-organised monastic collection comprising around 1,750–1,790 manuscripts, of which approximately 1,100–1,150 survive today, notably at the Médiathèque du Grand Troyes. The manuscripts covered a wide range of subjects, including biblical, patristic, theological, liturgical and classical texts, and were arranged according to a classification system and shelf marks that were refined throughout the fifteenth century. This reflects a sustained and sophisticated approach to managing the library.

#### Manuscripts in Cambridge libraries

##### Cambridge University Library

- **MS Kk.1.24**. This is a fragmentary gospel book (see https://cudl.lib.cam.ac.uk/view/MS-KK-00001-00024/1), containing substantial sections of the gospels of Luke and John, but lacking any large scale decoration, as is often found in Insular gospelbooks, other than the ornamentation of minor initials. Palaeographical analysis of the half-uncial script, suggests that the codex was made in the kingdom of Northumbria, probably in the first half of the eighth century (c. 700–750 CE). In some places, rubrics added between the lines indicate the passages that were to be read aloud on feast days, such as the Nativity of Mary; in other places letters indicate the sections of Passion narrative that were to be performed during the liturgy at Easter time. Another leaf from this manuscript is now divided between London, British Library, Cotton MS Tiberius B.v, fol. 76 and London, British Library, Sloane MS 1044, fol. 2.
- **MS Ll.1.10**. The Book of Cerne (see https://cudl.lib.cam.ac.uk/view/MS-LL-00001-00010/1) is so named after its later medieval provenance and subsequent additions made at Cerne Abbey in Dorset, UK. It is a Latin prayerbook that was probably made in the Anglo-Saxon kingdom of Mercia. Internal textual evidence suggests that it was made as the personal prayerbook for Bishop Æthelwald of Lichfield (818–830 CE). As such, it is one of only a small number of Anglo-Saxon manuscripts that can be securely dated to the ninth century.

##### The Parker Library, Corpus Christi College

- **MS 144**. This manuscript is known as the Corpus Glossary (see https://parker.stanford.edu/parker/catalog/mz111xq7301). It was made in southern England, probably Canterbury, Kent, in the first half of the ninth century, probably c. 800. The text contains two glossaries of Latin words, the first arranged in alphabetical order, and the second in AB order (by the first two letters of each word). Most words are provided with a Latin synonym, but some are provided with a synonym in Old English. Glossaries of this type are also found on the Continent, in contexts where people were learning Latin, and are often Insular in origin. Sample CL299 is from this part of the manuscript. At the back of the volume, two flyleaves are from a twelfth-century manuscript in Irish script, containing a fragment of a treatise on grammar by Priscian. Sample number CL301 is from these flyleaves.
- **MS 199**. This manuscript was written by a Welsh scribe named, Ieuan ap Sulien (d. 1137 CE; see https://parker.stanford.edu/parker/catalog/sk095st1718), who added some of his own verses in Welsh to its margins. It was made *c*. 1090 CE at Llanbadarn Fawr, near Aberystwyth in West Wales. The text is a copy of Augustine’s *De trinitate*.
- **MS 272**. This manuscript is known as the Achadeus Psalter (see https://parker.stanford.edu/parker/catalog/gv751fq0828). It contains (alongside the psalms) canticles, a litany, and prayers. On fol. 150r is an inscription, in gold uncials, saying that Count Achadeus ordered the book to be made (ie: paid for it). Achadeus is a Frankish name. The litany starts on fol. 151r and requests prayers for Pope Marinus (882–March 884 CE), King Carloman (of the West Frankish kingdom) (882–4 CE), and Fulk, archbishop of Reims (consecrated March 883–900 CE); these combine to give a date for the copying of the litany between March 883 and May 884 CE, and links at least that part of the manuscript to Reims. The litany is richly decorated with architectural arcades in wholly Frankish style. Marginal additions (from Cassiodorus’s commentary on the Psalms) show that the manuscript was in England by the eleventh century, and was in Canterbury in the later middle ages.

##### Wren Library, Trinity College

- **MS B.10.4**. The Trinity Gospels is one of the most lavishly produced Gospel Books to survive from late Anglo-Saxon England. Datable from script and decoration to the early decades of the eleventh century, it belongs to a small group of ‘deluxe’ Gospel Books whose closely related illumination styles, scripts, and parchments point to elite centres of manuscript production in southern England. Rich pigments and extensive gold attest to the considerable expense and technical skill involved in the production of the Trinity Gospels. The manuscript features heavily gilded canon tables, as well as gilded initials throughout. It is also the only surviving late Anglo-Saxon Gospel Book which preserves double-page illuminated openings preceding each of the four Gospels, complete with full evangelist portraits and decorated text. Although the manuscript cannot be pinpointed to a single place of origin, its script, a fine Anglo-Caroline minuscule, was surely written by a single, highly skilled scribe associated with the period’s finest English scribes. The scribe’s hand has been identified in other manuscripts also of uncertain Southern English origin but which share characteristics with manuscripts of the same date for which Christ Church, Canterbury, and Peterborough Abbey have been suggested as possible places of origin. Analysis of the parchment suggests a high degree of intentionality in its manufacture. In an age when the word for wealth *(feoh)* also meant cattle, a decision was made to construct the Trinity Gospels entirely from calfskin. Most folios are full, unblemished skins, indicating the deliberate selection and preparation of exceptionally small and fine hides. By contrast, the heavily illuminated pages appear on significantly thicker —but similarly unblemished—parchments likely made from older calfskins, a material choice that would have been necessary both to prevent show-through and support the physical weight of the pigments and gold. While the Trinity Gospels remain remarkably well preserved overall, most pages show evidence of mould damage and the early twentieth-century conservation treatment undertaken to address it.
- **MS B.10.5**. The origin of this manuscript is disputed, with either Ireland or Northumbria proposed as possible places of origin. The script dates it to the first half of the eighth century (c. 700–750 CE). It contains copies of the Epistles of Paul (see https://mss-cat.trin.cam.ac.uk/Manuscript/B.10.5), and has many marginal glosses. Some leaves of the manuscript are now kept separately, in London, British Library, Cotton MS Vitellius C. viii, fols. 85–90.

##### Vienna, Österreichische Nationalbibliothek

- **Cod. 1224** (see https://onb.digital/result/13226B0F). This is a copy of the four gospels, written by a named scribe, Cutbercht, who wrote a good insular half uncial script, alongside insular minuscule and uncial, and was working at a scriptorium in Francia, probably Salzburg (Austria), in the later eighth century (c. 785-790 CE). A note on fol. Ov records ‘ *Cutberc\h/t. scripsit ista. IIII. evangelia*’ (‘Cutbercht wrote these 4 gospels’), and Cutbercht’s hand is found in other manuscripts of similar date linked to Salzburg. The manuscript has evangelist portrait pages and large insular style initials at the start of each gospel text.

##### “Le Bouteiller”, Louvres, France

The samples RF002 to RF009 stem from an archaeological assessment that was carried out in 2012 by the INRAP under the supervision of F. Gentili in Louvres, France, about 25 kilometers north of Paris, to evaluate the archaeological potential of what used to be the vegetable garden of a farm established in the 16th century (*92*). A bowl-shaped pit containing nine articulated sheep skeletons was brought to light; no other noteworthy features were identified by the survey, and no excavations of the plot were mandated. The sheep were radiocarbon-dated to the Early Modern period (1658-1888 cal. CE, 175±30 BP, Poz-85860, 2σ calibration), and determined through archaeological and paleopathological evidence to have died in a single mass mortality event(*23*). Subsequent palaeogenetic analyses confirmed the animals had died from an outbreak of sheeppox, a disease common in the area at the time (*27*).

#### Lambeth Palace Library

The Lambeth Bible (Lambeth Palace Library MS 3) is one of approximately a dozen surviving giant Romanesque Bibles produced in England. Alongside the Winchester and Bury Bibles, it is regarded as among the most finely illuminated, including six full-page paintings (one of which features on fol. 6r) and twenty-four historiated initials. The first volume, containing the text of Genesis to Job, is held at Lambeth Palace Library while the second is in Maidstone Museum & Art Gallery (MS P.5). MS 3 is believed to have received its current binding in the fifteenth or sixteenth century, at which time the pastedown on the backboard (from which sample LambethBible3MSXXXXXPastedown1rightXXXXXimg330 was taken) was likely added.

#### Veterinary pathology museum specimen, Vienna

The specimen from the Veterinary pathology museum at the Vetmeduni Vienna shows irregular confluent papules and nodules with clear demarcation lines. The skin is sparsely haired or alopecic (Figure S20A). Histologically the nodules prove to be full-thickness dermal necroses, demarcated by inflammatory infiltrates (Figure S20B). In the necrotic tissue the epithelium of the epidermis and hair follicle infundibula shows severe parakeratotic hyperkeratosis and obvious ballooning degeneration especially at the stratum spinosum (Figure S20C). Small pustules are present and the necrotic tissue shows bacterial overgrowth. The necrotic superficial dermis is hyperaemic, oedematous and moderately infiltrated by leucocytes. The thickened subcutis underneath the necrosis is likewise hyperaemic and shows moderate perivascular to diffuse mixed inflammatory infiltrates. Unequivocal sheppox cells with clear cytoplasm and marginated chromatin, eosinophilic inclusion bodies, vasculitis or intravascular thrombi are not visible.

The gross and microscopic lesions of the specimen correspond to typical pathological lesions caused by sheeppox virus as described by Plowright et al., 2012 (*93*). In their experiments between the 11th and 14th days after inoculation of sheeppox virus the surface and follicular epithelium was necrotic with swelling and vacuolation of the epidermis and infundibular epithelium. The whole dermis was necrotic and in contrast to earlier stages there were very few sheeppox cells visible, which meets the changes seen in the museum sample. The original protocol no. 252a from the 9th of March 1917 states: “skin pieces of a sheep; Wrana (Böhmen); sheeppox”. Wrana is the German name of Vraný, Czech Republic and was part of the Austro-Hungarian Empire at that time. Organs from diseased domesticated animals from the whole empire regularly were sent to the Tierärztliche Hochschule at Vienna for examination in the early 20th century.

### Sampling and extraction

Tooth samples: Sample MYRZ180 was processed in a dedicated ancient DNA laboratory in Trinity College Dublin, Dublin. All work surfaces were routinely cleaned with dilute sodium hypochlorite; researchers were full-body PPE at all times, with frequent changing of gloves. The tooth was sectioned along the cementoenamel junction, and the root was further halved. Material from the pulp chamber and the interior of the root was collected using a dental drill, yielding 56 mg of powder. A single tooth specimen (RUB002) was prepared at the Christian Albrecht University of Kiel in a restricted-access, decontaminated laboratory using strict ancient DNA protocols. Work surfaces, instruments, reagents, and containers were routinely decontaminated with 7.5% bleach, DNA-free water, and UVC irradiation as appropriate. The sample was first decontaminated by UVC exposure, followed by immersion in 7.5% sodium hypochlorite with agitation, then subjected to three sequential washes in laboratory-grade DNA-free water before being fully dried in a desiccator for a minimum of four days. Prior to powdering, the sample and associated consumables were again UVC irradiated, and all milling equipment was thoroughly decontaminated using bleach, DNA-free water, isopropyl alcohol, and UVC exposure between samples. The dried sample was entirely pulverized in decontaminated stainless steel mill jars with grinding balls using a mixer mill, with repeated cycles as needed until fully powdered, after which the resulting powder was transferred into a labeled 2 mL tube and stored at room temperature. Extraction was performed on a ∼80mg subsample for RUB002.

For DNA extraction, both teeth samples were processed in a dedicated ancient DNA laboratory in Trinity College Dublin, Dublin. Sample RUB002 was subject to the extraction protocol described in Mattiangeli et al., 2023 (*94*). Briefly, a dilute sodium hypochlorite wash (0.5% over 15 min, followed by three 1 ml H_2_O washes) was performed. Following this, the sample was subject to a 1 ml 0.5M EDTA pre-digestion at 37 °C over 30 minutes, with frequent vortexing, followed by removal of the EDTA supernatant. The remaining pellet was digested overnight using 1 ml of an extraction buffer (per sample: 17 μl N-Laurylsarcosine, 20 μl 1M Tris-HCl, 940 μl 0.5M EDTA, 13 μl 50 U/ml proteinase K [UVing the mastermix for 30 min prior to proteinase K addition]), at 37 °C with frequent agitation. After tube spin down (17,000 g for 10 min), the resulting supernatant was purified using Roche High Pure Extender Assembly columns and 13 ml of modified PB buffer (per sample: 0.42 ml 3M sodium acetate, 0.33 ml 5M sodium chloride, 12.25 ml PB buffer (Qiagen). DNA was eluted in 50 μl EBT. Sample MYRZ180 was extracted following the same protocol, except that the sodium hypochlorite wash was omitted and the EDTA pre-digestion step was reduced to 15 minutes.

Parchment: DNA was extracted from the parchment eraser samples following the protocol of Teasdale et al., 2017 (*38*) based on Gamba et al., 2014 (*95*). Briefly, the parchment sample was incubated at 55 °C for 24 h in 0.25 mL of lysis buffer [1 M Tris·Cl (pH 7.4), proteinase K, 10% SDS, 0.5 M EDTA]. Following this incubation, the sample was centrifuged at 16,249 × g for 1 min to precipitate any undigested material; the supernatant was then removed, and DNA was recovered using the Qiagen Minelute kit. The first DNA binding step of this protocol (Qiagen Minelute) was repeated two times due to the large volume of starting material, and a final elution volume of 20 μL was used.

Pathology collection: We performed extractions in a dedicated ancient DNA laboratory at the Helmholtz Institute for One Health (Greifswald, Germany). Small tissue samples (ca 25mg) were washed twice in PBS before being placed in tubes containing 1.4mm ceramic beads and 180 µL of the buffer ATL from the DNeasy Blood and Tissue kit (Qiagen). The tubes were heated 15 min at 98°C to reduce molecular cross-links induced by formalin fixation, before being placed in a bead beater for homogenization. After cooling, we added proteinase K to the homogenate and incubated at 56°C for one hour. After digestion, the lysate was centrifuged at 20,000 g for 1 min to pellet remaining tissue debris. The supernatant was transferred to a fresh tube and used for extraction according to manufacturer’s instructions with the DNeasy Blood and Tissue kit. DNA was eluted in a 100 µL buffer AE.

### Library building and shotgun sequencing

Teeth: Double-stranded DNA libraries were prepared following the protocol of Meyer and Kircher (*96*). For each sample, one library was constructed without uracil-DNA glycosylase (UDG) treatment using 12–16 μl of input DNA to preserve authentic damage patterns. All remaining libraries were treated with 5 μl UDG (New England Biolabs, M5505L) at 37 °C for 1 h prior to library construction. Libraries were dual-indexed and PCR-amplified for 12 cycles, then shotgun sequenced on an Illumina NovaSeq X platform with 150 bp paired-end reads. Sequencing statistics and metadata for each indexed amplification are reported in Table S2.

Parchment: Illumina sequencing libraries were produced for each of the samples and appropriate controls following the protocols of Teasdale et al., 2017 (38) and Gamba et al., 2014 (95). Sequencing of multiplexed pools was performed on an Illumina NovaSeq 6000 S4 platform (PE 200bp) at the GeoGenetics Sequencing Core, Copenhagen. ssDNA libraries for samples TCCL03 to TCCL10 were prepared following Troll et al., 2019 (*97*) and using Santa Cruz Reaction method developed by Kapp et al., 2021 (*98*). DNA sequencing was performed on an Illumina NovaSeq X platform using 150 bp PE reads.

Pathology collection: We prepared Illumina sequencing libraries from dsDNA using the NEBNext Ultra II DNA Library Prep kit for Illumina and unique dual indices (New England Biolabs). We shotgun-sequenced them on an Illumina MiniSeq platform with 75 bp paired-end reads.

### Radiocarbon dating

A small amount of specimen powder (100-1000 mg) of samples MYRZ180 and RUB002 were subject to radiocarbon dating, by CIRAM (Martillac) and Oxford Radiocarbon Accelerator Unit (Oxford) respectively. The resulting uncalibrated ages (CIRAM-8056: 3021 ± 30; OxA-45742: 3440 ± 19) were calibrated using OxCal4 (*99*) and IntCal20 (*100*). Calibrated 2σ ages for both samples, and those whose radiocarbon age was produced previously, are presented in Table S1. CIRAM-8056 was produced via cold HCl etching and gelatinization via hot acid extraction; no ultrafiltration was performed. OxA-45742 was produced with the following pre-treatments: NaOH rinse, gelatinization, ultrafiltration.

### Screening of sequencing reads

One fastq file per sample from available ancient ruminant datasets (PRJEB26011, PRJEB31621, PRJEB40573, PRJEB43881, PRJEB75678, PRJEB61808, PRJEB59481, PRJEB75467, PRJEB63473,PRJEB69690,PRJEB81145,PRJEB78495) and 35 newly reported material were screened. Following adapter removal, host-derived reads were removed by aligning reads with Bowtie2 against a concatenated reference genome comprising sheep (ARS-UI_Ramb_v2.0), goat (ASM170441v1), and cattle (ARS-UCD1.2). No mapping quality filter was applied. Reads that did not align to the host reference were retained for downstream analyses.

Non-host reads from all screening libraries were merged, and metagenomic profiling was performed using KrakenUniq v1.0.4 with a custom database containing a microbial version of NCBI nt combined with human and complete eukaryotic reference genomes. KrakenUniq reports both read counts and unique k-mer counts for each taxon, enabling sensitive detection of low-abundance taxa and estimation of coverage breadth.

To discriminate true from spurious taxonomic assignments, we applied the scoring framework developed by Guellil et al. (*101*), which computes an E value defined as *E* = (K/R) × C, where *K* is the number of unique k-mers, *R* is the number of assigned reads, and *C* is the estimated breadth of coverage. E values were calculated for each KrakenUniq assignment, and results were filtered at the species level for Goatpox virus, Sheeppox virus, and Lumpy skin disease virus. Assignments were retained only if supported by at least five reads and an E value ≥ 0.001. Because Capripoxvirus genomes are highly conserved, KrakenUniq systematically assigned a subset of reads to multiple Capripoxvirus species. To resolve species identity, we identified the Capripoxvirus species exhibiting the highest E value among Capripoxvirus assignments for each sample and it was systematically SPPV (Figure S1, Table S1). Following alignment to the SPPV reference genome using the same parameters described in the *Alignment on Sheeppox virus genome* section (below), the edit distance between sequencing reads and the reference was calculated using a custom Python script.

### Host species mtDNA assessment

Host species identification was performed by analyzing mitochondrial DNA, for SPPV-positive samples with a SPPV genome ≥ 0.1× (i.e. those indicated in Table 1). Sequencing reads were aligned to a concatenated reference of complete mtDNA genomes from goat (NC_005044.2), sheep (NC_001941.1), and cattle (V00654.1) using the bwa aln algorithm with relaxed parameters (-l 1024 -n 0.01 -o 2). Alignments were filtered to retain reads with a minimum mapping quality of 30, and PCR duplicates were removed (Table S3). Samples with > 0.9 of aligned reads mapping to a single species were assigned to that species, whereas samples with < 0.9 were classified as mixed, with the majority species identified together with any minor species representing ≥ 0.1 of the aligned reads. Species assignments are reported in Tables 1 and S1. For each sample, mitochondrial consensus sequences were generated using ANGSD (*102*) (-doFasta 2). For samples showing evidence of mixed species, multiple consensus sequences were generated from the competitive alignment BAM file. The resulting mitochondrial consensus sequences were aligned using MAFFT (parameters: --adjustdirection --maxiterate 1000) (*103*), and a maximum likelihood phylogeny was inferred with IQ-TREE2 (*104*) using the TIM2+F+I+R3 nucleotide substitution model selected by ModelFinder and 1,000 ultrafast bootstrap replicates. Phylogenetic placement of the mtDNA sequences was used to confirm species and haplogroup assignments (Figure S2). Observed haplogroups were consistent with those expected for each host species, including in samples in which mtDNA from a given species was present only as a minor fraction of the reads.

**Table 1:** Metadata of SPPV-positive samples over 0.1× mean genome coverage. ^1^ indicates 95% calibrated range of ^14^C-dated samples. ^2^ indicates samples with possible mixed species signals; species most represented in competitive alignment is presented here (see Table S3 for details). ^3^ indicates that the page was replaced at a later date and the sample age is potentially later than indicated (see Supplementary Note 1). ^4^ indicates likely identical sequences, see (*27*).

| Sample | Material | Age | Context | Host | Origin | Coverage |
| --- | --- | --- | --- | --- | --- | --- |
| MYRZ1<br>80 | Tooth | 1875-1645<br>BCE<sup>1</sup> | Bronze<br>Age | Sheep | Myrzhik; Kazakhstan | 260.57 |
| RUB002 | Tooth | 1336-1130<br>BCE<sup>1</sup> | Bronze<br>Age | Sheep | Rublevo-VI; Russia | 86.52 |
| TCCL02 | Parchment | 700-750 CE | Medieval | Cattle | *Pauline Epistles*; Ireland or<br>Northumbria, UK | 50.51 |
| CL299 | Parchment | 800-850 CE | Medieval | Cattle<sup>2</sup> | *The Corpus Glossary*; probably<br>Kent, UK | 212.98 |
| YG15 | Parchment | 990-1020 CE | Medieval | Cattle | *The York Gospels*; Canterbury, UK | 1.81 |
| YG16 | Parchment | 990-1020 CE | Medieval | Cattle | *The York Gospels*; Canterbury, UK | 10.81 |
| PEM01 | Parchment | 1100-1300 CE | Medieval | Cattle<sup>2</sup> | *XI 249*, St. Florian monastery; Austria | 0.31 |
| PEM02 | Parchment | 1000-1100 CE | Medieval | Cattle | *XI 249*, St. Florian monastery; Austria | 0.18 |
| PEM03 | Parchment | 1000-1100 CE | Medieval | Cattle<sup>2</sup> | *XI 249*, St. Florian monastery; Austria | 0.58 |
| PEM04 | Parchment | 1180-1200 CE | Medieval | Cattle | *III 1*, St. Florian monastery; Austria | 0.35 |
| PEM05 | Parchment | 1180-1200 CE | Medieval | Cattle<sup>2</sup> | *III 1*, St. Florian monastery; Austria | 0.18 |
| TP04 | Parchment | 1201-1300 CE | Medieval | Sheep | Clairvaux Monastery; France | 1.60 |
| TP02 | Parchment | 1301-1400 CE | Medieval | Cattle<sup>2</sup> | Clairvaux Monastery; France | 0.35 |
| YG01 | Parchment | 1300-1600 CE | Medieval | Sheep<sup>2</sup> | Addition to *The York Gospels*; probably Yorkshire, UK | 1.10 |
| BA27 | Parchment | 1380 CE | Medieval | Sheep | *MOR 3*; Bossall, Yorkshire, UK | 8.17 |
| 16R | Parchment | 1420-1460 CE | Medieval | Goat<sup>2</sup> | *Speculum historiale*; Upper Rhine, Switzerland/Germany | 0.76 |
| 79RR | Parchment | 1420-1460 CE<sup>3</sup> | Medieval | Goat<sup>2</sup> | *Speculum historiale*; Upper Rhine, Switzerland/Germany | 5.20 |
| 80R | Parchment | 1420-1460 CE | Medieval | Goat | *Speculum historiale*; Upper Rhine, Switzerland/Germany | 1.12 |
| 48R | Parchment | 1420-1460 CE | Medieval | Goat | *Speculum historiale*; Upper Rhine, Switzerland/Germany | 0.14 |
| TP01 | Paper | 1450-1455 CE | Medieval | Cattle<sup>2</sup> | Clairvaux Monastery; France | 0.23 |
| RF002<sup>4</sup> | Tooth | 1658-1888 CE<sup>1</sup> | Early Modern | Sheep | Louvres, France | 12.09 |
| RF003<sup>4</sup> | Tooth | 1658-1888 CE<sup>1</sup> | Early Modern | Sheep | Louvres, France | 262.93 |
| RF009<sup>4</sup> | Tooth | 1658-1888 CE<sup>1</sup> | Early Modern | Sheep | Louvres, France | 1.55 |
| ANC255 | Veterinary Pathological Museum | 1917 CE | Early Modern | Sheep | Vraný, Czech Republic | 110.14 |

### *Capripoxvirus* capture design

The design of the *Capripoxvirus* capture probe was previously reported (*27*). A set of custom in-solution 3x-tiled 80bp RNA baits (myBaits®) were designed and synthesized by Daicel Arbor Bioscience, targeting *Capripoxvirus* DNA, as initially reported in (*27*). The bait design was based on reference grenomes sequences for *Capripoxvirus* members (Sheeppox virus: NC_004002.1; Goatpox virus: NC_004003.1; Lumpy Skin Disease Virus, NC_003027.1), with low complexity regions masked. Baits matching to host livestock genomes (Sheep: GCF_016772045.2; Goat: GCF_001704415.2_ARS1.2; Cattle: GCF_002263795.3_ARS-UCD2.0) were excluded (5 baits), as were baits with more than 35% repeat content. The final set of 80bp baits amount to 15,646 unique probes, designed to enrich for DNA across *Capriopoxvirus*.

### *Capripoxvirus* capture application

Amplified, dual index-incorporated libraries were targeted for enrichment for SPPV DNA, through application of the RNA baits described above. Amplified libraries were combined into separate pools (samples with a similar viral DNA representation were pooled together), with the aim of pooling at 200–2000 ng of total DNA when possible, and combining equal amounts of DNA per sample. Pools were desiccated and worked back up to 7 μl of lab grade H2O. Pools were then subject to overnight (∼20 hours) capture using the RNA baits, following the manufacturer’s instructions for degraded DNA (“High Sensitivity”), including setting the annealing temperature to 55°C and performing a second round of annealing and amplification (8 cycles) after the first amplification (16 cycles). In some cases, only one round of capture/annealing was performed; the number of capture rounds per sample is indicated in Table S1. Amplification was performed using KAPA HiFI Hotstart (Roche). Purified, captured libraries were then subject to shotgun sequencing for 150bp paired end reads on NovaSeq X platforms (Macrogen Europe, Amsterdam). Due to the high representation of SPPV DNA in MYRZ180, this sample was subject to deep shotgun sequencing only. RUB002 was subject to both deep shotgun sequencing and targeted enrichment. We also note that the initial screening of parchment samples reported here in many cases exceeds 100 million reads per sample, prior to targeted enrichment. Previously published data from two petrous samples from Neolithic Serbia and Bronze Age Uzbekistan (Blagotin 17 and Bulak2) each had a single read detected for Capripoxvirus. Both samples were subjected to capture, but failed (i.e. no reads assigned to Capripoxvirus or aligned SPPV). We reported them (Table S1), but we do not consider these true-positive SPPV detections.

### Alignment on Sheeppox virus genome

Captured and shotgun reads were trimmed and host-filtered using the same parameters as described above. The resulting screened data and captured reads were then aligned using the bwa aln algorithm with relaxed parameters (-l 1024 -n 0.01 -o 2) to a modified version of the Sheeppox virus (SPPV) reference genome (NC_004002.1). Because the SPPV genome contains two identical inverted terminal repeat (ITR) regions at its extremities, reads originating from these regions map ambiguously to both ends of the genome, resulting in a mapping quality score of zero and subsequent exclusion from downstream analyses. To prevent this issue, we removed the terminal 2,213 bp from the reference genome using seqkit (*105*), thereby enabling unique mapping of reads to the first extremity. Reads with mapping quality under 30 were filtered out with samtools v1.19.2, duplicates were removed and bam coming from the same samples were merged with picard MarkDuplicates and MergeSamFiles v2.26.11. To confirm that both capture-enriched and shotgun sequencing reads originated from SPPV, the same alignment procedure was applied to the LSDV (NC_003027.1) and GTPV (NC_004003.1) reference genomes. Pairwise genetic distances between the aligned reads and each reference genome were then calculated using a custom Python script. Ancient DNA typical damage pattern was then assessed using DamageProfiler v1.1 (*106*). Reads in the merged BAM files were trimmed by 4 bp at both 5′ and 3′ ends using BamUtil v1.0.15 (*107*).

### Localized sampling of TCCL02

Sampling: To determine the localization of SPPV DNA on a particular parchment page, we performed sampling on a single bifolium (single piece of skin), across both folios (pages), on either the flesh side or hair side. In this case, we sampled folios 55–56 from the Pauline Epistles (Cambridge, Trinity College, MS B.10.5), sampling both right (r = recto, right) and left pages (v = verso, left): 55r and 56v = hairside; 55v and 56r = flesh side.

We obtained six samples using eraser-based sampling. Pairs of samples were taken from the same location, with the initial sampling aim to obtain the most topical DNA. TCCL03 and 04 were sampled from the lower margin of fol. 55r (hairside). TCCL05 and 06 were taken from the verso of fol. 55 (flesh side), from the lower margin, towards the spine. TCCL07 and 08 were taken from the outer margin of fol. 56r (flesh side), and should map to TCCL05 and TCCL06. Finally, we performed fibre sampling: TCCL09 was two fibres taken from fol. 56r (flesh side), lower margin, from the surface; TCCL10 were two fibres from fol. 55r (hairside), upper margin, from the surface.

DNA extraction and library creation: Samples were extracted following (*94*), excluding the dilute bleach and EDTA pre-washes. DNA fragments were converted into ssDNA libraries using the Santa Cruz approach (*98*). The resulting libraries were sequenced 150bp paired-end using an Illumina NovaSeq X platform (Macrogen Europe, Amsterdam).

Data generation and analysis: Data were processed following the same workflow described in the “Screening of sequencing reads” section. Non-host reads were then aligned to the modified Sheeppox virus reference genome (NC_004002.1) using BWA, as described in the “Alignment on Sheeppox virus genome” section. For subsequent analysis, SPPV aligned reads (after mapping quality 30 and duplicate removal) were normalized to 20 million reads. A Wilcoxon rank-sum test was computed to assess statistically significant differences between hair and flesh sampling sites; comparison of the distributions are shown in Figure S3. All statistical analyses were conducted using the *tidyverse, ggpubr* and *rstatix* packages in R.

### Phylogenetic analysis

To assess the phylogenetic placement of ancient SPPV genomes within modern diversity, we constructed a maximum likelihood phylogeny including 16 complete wild (i.e. excluding passaged or vaccine strains) modern SPPV genomes with available collection dates from NCBI, excluding one genome with an excessive branch length (PQ014465.1), alongside ancient SPPV genomes with a mean sequencing coverage greater than 5× (*n* = 9). Ancient genomes with coverage between 0.1× and 5× and redundant sequence from(*108*) were subsequently placed onto this ML tree using pathPhynder v1.2.4 (*45*).

To generate reliable ancient sequences for ML inference, consensus sequences for samples with mean coverage > 5× were generated after variant calling with FreeBayes v1.3.6 (*109*). Minimum coverage thresholds were set to 3× for samples averaging 5-11× and 10× for those > 11×. FreeBayes was run using a minimum alternate allele fraction of 0.8, minimum mapping quality of 30, the --report-monomorphic flag to retain homozygous reference sites, ploidy set to 1, and a minimum alternate count equal to the coverage threshold to ensure robust SNP support. Variant calls were filtered with bcftools view v1.17, with sites having QUAL < 30 flagged as LOWQUAL. Variants were then normalized, split, and indels removed to retain SNP-only calls. LOWQUAL sites were set to missing, and consensus sequences were generated with bcftools consensus using the modified Sheeppox virus reference genome. The -a N -M N parameters were applied to mask uncovered and LOWQUAL positions with “N,” ensuring low-coverage or filtered sites were appropriately masked.

Then, consensus sequences of ancient samples, modern SPPV isolates genomes, Goatpox virus reference genome (NC_004003.1) and Lumpy skin disease virus reference genome (NC_003027.1) were aligned using MAFFT (parameters: --adjustdirection --maxiterate 1000) (*103*). The resulting alignment was used to infer a maximum likelihood phylogeny in IQ-TREE2 (*104*), employing the nucleotide substitution model TVM+F+I+G4 as determined by ModelFinder and 1,000 ultrafast bootstrap replicates. The phylogenetic tree was rooted with LSDV and GTPV.

Because pathPhynder requires a shared set of variant positions across samples, a core genome VCF was generated with ancient and modern genomes by aligning modern synthetic reads produced with ART (parameters -l 150 -f 50) (*110*) using the same alignment parameters described in the “*Alignment on Sheeppox virus genome*” section. Variants were called with bcftools mpileup v1.15, using a minimum mapping quality of 30 and base quality of 20. Variant calling was performed with bcftools call (*111*) with ploidy set to 1. Per-sample variant calls were filtered with bcftools view to retain only sites with a QUAL score ≥30, and a sequencing depth of ≥ 3. Finally, this core genome SNP dataset was used to project low-coverage ancient genomes onto the high-confidence ML reference tree, using pathPhynder (*45*) with default parameters.

To assess the evolutionary relationships among the capripoxvirus samples and nine related subfamily *Chordopoxvirinae* species, we aligned genomes with Mafft v7.453 (*103*) using automatic optimisation and default parameters. The region corresponding to LSDV027-LSDV123 was excised using LSDV genome KX894508.1 as a reference. The evolutionary relationships of this core genome dataset were reconstructed using RAxML-NG v1.2.0 (*112*) with 1,000 bootstraps and with a GTR model and gamma substitution rate heterogeneity, selected by modeltest-ng (*113*). The phylogeny was rooted using the nine non-capripoxvirus chordopox samples, and was visualised using ape v5.7-1 (*114*), ggtree v3.8.2 (*115*), phangorn v2.11.1 (*116*) and phytools v2.0-3 (*117*). We repeated this using a site-heterogeneous substitution model GTR+FO*H4 in IQ-TREE that included optimised base frequencies and four frequency classes to allow sites to have different composition rates to address potential long branch attraction effects stemming from the site-homogeneous RAxML model. This results in the same topological structure and nearly identical branch lengths.

### Verification of Temporal Signal

The SPPV dataset was tested for the presence of a temporal signal, an assumption required for molecular clock calibration. First, a maximum likelihood phylogeny was inferred using the same dataset described in the Phylogenetic analysis section, excluding LSDV and GTPV genomes, and rooted using MYRZ180, the oldest available genome. This tree was analyzed in TempEst (*47*) to assess the relationship between root-to-tip genetic distance and sampling time. Tip dates were assigned using the median of the historical age estimates for parchment-derived genomes and the median calibrated ages for radiocarbon-dated ancient sequences. A strong linear correlation between root-to-tip distance and sampling time was observed (R^2^ = 0.96, Figure S6), indicating a strong temporal signal. To further validate this result, the dataset was analyzed using the Bayesian Evaluation of Temporal Signal (BETS) (*46*) framework. Four models were implemented in BEAST v1.10.5 (*118*), combining either a strict or an uncorrelated lognormal molecular clock with the inclusion or exclusion of temporal information. The model incorporating an uncorrelated lognormal clock and temporal information showed the highest marginal likelihood (Figure S7) and was strongly favored over the corresponding model without temporal information (log Bayes factor = 4.34, Table S5), confirming the presence of a temporal signal consistent with the TempEst analysis. The analysis was subsequently repeated after excluding the Bronze Age genomes (MYRZ180 and RUB002) to assess temporal signal within the medieval-to-modern timeframe, and a positive correlation between root-to-tip genetic distance and sampling time was still observed (R^2^ = 0.68).

To assess temporal signals in the *Capripoxvirus* datasets, we analysed the full dataset using Clockor2 (*119*). A maximum likelihood phylogeny of capripoxvirus genomes was first midpoint-rooted; this; this indicated SPPV as the potential root, and the tree was subsequently rooted on SPPV. A weak global temporal signal was observed across the full *Capripoxvirus* dataset, as shown by the correlation between root-to-tip genetic distance and sampling time (R^2^ = 0.15; Figure S19), indicating limited clock-like signal. In contrast, temporal signal varied among individual Capripoxvirus species when analysed separately using Clockor. SPPV exhibited a strong temporal signal (R^2^ = 0.97), whereas LSDV showed a much weaker signal (R^2^ = 0.16). Due to the heterogeneity in clade branch lengths in GTPV compared with LSDV and SPPV, species-specific Clockor analyses were performed on individual GTPV clades rather than on the full GTPV dataset. All analysed clades displayed no temporal signal (clade 2.1: R^2^ = 0.06; clade 2.2: R^2^ = 0.00; clade 2.3: R^2^ = 0.003).

### Time to Most Recent Common Ancestor (tMRCA) of SPPV sequences

To investigate the tMRCA of all SPPV sequences and the divergence times of subclades, we constructed time-calibrated Bayesian phylogenies using BEAST v1.10.5 (*48*). Full genomes of ancient samples with at least 5× coverage, together with 16 modern genomes, were aligned using MAFFT (-adjustdirection --maxiterate 1000) (*103*).The optimal nucleotide substitution model was then determined with ModelFinder (model:TVM+F+I). For ancient samples with radiocarbon dates, the probability density function of the C_14_ calibration was used as a prior for tip dates via the empirical calibrated radiocarbon sampler (ECRS) (*120*). For manuscript samples which are lacking radiocarbon dates, a uniform prior based on estimated date ranges from historical records was applied. Beast was run under a GTR+T+I substitution model, with a constant-size coalescent tree prior and an uncorrelated lognormal (UCLD) relaxed molecular clock model (*51*), given the results of the BETS test (Figure S7, Table S5). Three independent Markov Chain Monte Carlo (MCMC) chains of 250 million steps each were run. Convergence and effective sample sizes (ESS) were assessed in Tracer v1.7.2 using 20% burn-in, confirming excellent mixing (ESS >> 300) and convergence between runs. The resulting tree files were combined with LogCombiner, applying 20% burn-in, and maximum clade credibility (MCC) trees were generated with TreeAnnotator using 0% burn-in, as burn-in had already been removed during combination.

### Tip-dating of manuscript samples

To evaluate whether the SPPV molecular clock could be used to tip-date parchment-derived samples of an unknown age with a minimum sequencing depth of 5×, two time-calibrated Bayesian phylogenetic model sets were tested. In the first model set, Bayesian phylogenetic analyses were performed using the same dataset and tip-date priors as described in the *Time to Most Recent Common Ancestor (tMRCA) of SPPV sequences* section, except that the sampling date of one of the five parchment-derived genomes was replaced by a uniform prior spanning the temporal range of the tree (from 0 to the median age of MYRZ180) i.e. five models in total, each including one parchment sequence with wide uniform prior. In the second model set, we explored five different datasets; in each we removed all parchment-derived genomes except one, which was assigned a uniform prior spanning the temporal range of the tree. That is, we attempted to estimate the age of the parchment-derived sequences using high quality, non parchment-derived sequences.

### Time-calibrated phylogeny of *Capripoxviruses*, recombination inference, and root estimation

To estimate the time to the most recent common ancestor of the *Capripoxvirus* genus and reconstruct its evolutionary history, we analyzed a set of capripoxvirus genomes using time-calibrated phylogenetic inference. This analysis included recombination-free wild-type LSDV genomes (n = 32) (*16*), wild-type GTPV genomes (*n* = 9), 15 modern SPPV genomes, and consensus sequences from ancient SPPV genomes with a minimum sequencing coverage of 5× (see Table S7 for sample details). Sequences were aligned using MAFFT with parameters described previously. Recombination has been shown to drive evolution in modern Capripoxviruses, notably between vaccine-derived LSDV strains and wild field strains (*121, 122*). To avoid phylogenetic biases caused by recombination, the alignment was analyzed with RDP5 (*123*) using seven complementary detection methods: RDP, GENECONV, BootScan, MaxChi, Chimaera, SiScan, and 3Seq. All analyses were run under default settings, with a Bonferroni-corrected P-value cutoff of 0.05, and only recombination signals supported by at least four of the seven methods were conserved. Regions of the alignment suspected of recombination were subsequently removed, resulting in a total of 4.16 kb removed from the alignment for a final alignment of 148,302 bp size.

A maximum likelihood tree was built from the filtered alignment to ensure the correct placement of each tip within its respective species, using IQ-TREE 2 with the nucleotide substitution model K3Pu+F+I+G4 as determined by ModelFinder, and 1,000 ultrafast bootstrap replicates (Figure S9).

The filtered alignment was then used for the construction of time-calibrated Bayesian phylogenies. For ancient samples with radiocarbon dates, the probability density function of the C_14_ calibration was used as a prior for tip dates via the ECRS (*120*). For manuscript samples which are lacking radiocarbon dates, a uniform prior based on historian-estimated date ranges was applied. Two independent Beast models were run under a GTR+T+I substitution model, with a constant-size coalescent tree prior and two different clock models, the UCLD relaxed molecular clock model and Shrinkage-based Random Local Clocks (*51, 52*). Because the UCLD model assumes independently distributed branch-specific rates, it may be suboptimal for evolutionary scenarios with structured or punctuated rate variation (*124*). We therefore also evaluated the shrinkage-based random local clock model (*52*), which uses Bayesian shrinkage priors to model heritable, structured rate shifts among branches, which may better fit our dataset of three *Capripoxvirus* lineages. Three independent MCMC chains of 300 million steps each were run. RootAnnotator (*50*) was then employed to count root positions across all trees in the posterior sample generated by the Beast runs. After analyzing the runs without any phylogenetic constraints, RPPs for the branches leading to the LSDV and GTPV clades were effectively close to zero (RPPs: Uncorrelated lognormal molecular clock model: LSDV: 0.005, GTPV: 0.001; Shrinkage-based Random Local Clocks: LSDV: 0.02, GTPV: 0.0002) whereas the branch leading to the SPPV lineage showed elevated support (approximately 0.44-0.93 under both clock models). In these unconstrained analyses, MYRZ180 and RUB002 were occasionally placed as the root of the *Capripoxvirus* phylogeny in a minor fraction of posterior trees (RPPs: Uncorrelated lognormal molecular clock model: MYRZ180: 0.3, RUB002: 0.003 MYRZ180&RUB002 together: 0.02; Shrinkage-based Random Local Clocks: MYRZ180: 0.02, RUB002: 0.0004 MYRZ180&RUB002 together: 0), which reduced the overall support assigned to the true SPPV root position and consequently produced a more recent (closer to the present) estimated split with a wider 95% HPD interval. Because maximum likelihood analyses clearly identify MYRZ180 and RUB002 as SPPV strains, we subsequently enforced monophyly of the three species groups (SPPV, GTPV, and LSDV). The analysis was subsequently repeated with an additional constraint enforcing the monophyly of samples according to their species assignment, by defining taxon sets in BEAST based on species labels and their maximum likelihood placement. Under these constraints, the branch leading to SPPV was unambiguously inferred as the root of the *Capripoxvirus* clade, receiving an RPP of ∼1.0 under both clock models (Figures 3a, S10, S14, S15 and Table S6). Additionally, to mirror the phylogenetic analyses performed on *Capripoxvirus* core genes, a time-calibrated Bayesian phylogeny was inferred using a UCLD relaxed molecular clock model, with rooting enforced on LSDV (Figure S16).

### Sliding window pairwise distances

To assess similarities between capripoxvirus species, pairwise distance calculations were performed on the initial *Capripoxvirus* MSA. Samples with coverage below 50× (YG16, BA27, and 79RR) were excluded to avoid the excessive presence of ambiguous bases (Ns), which could bias pairwise estimates. Regions identified as recombinant were masked, and the first and last 3,000 bp of the alignment were removed to prevent misalignment artifacts resulting from the absence of one inverted terminal repeat (ITR) in ancient samples and size variation in the ITRs across genomes. Global pairwise distances were first computed between all samples, and mean values were summarized at the species level. The same analysis was then repeated using a sliding-window approach across the alignment, employing 500 bp windows with a 100 bp step size; distributions are shown in Figure S11.

### De-novo assembly of SPPV genomes

De novo assembly was performed to accurately detect insertions and deletions, as poxviruses frequently contain indels that can cause frameshifts and inactivate genes. To assemble our ancient dataset with high accuracy, SPAdes v3.15.5 (*125*) was used in careful mode, which requires paired-end reads. For samples with a minimum coverage of 1×, AdapterRemoval was rerun without the --collapse option to retain only paired reads. These paired reads were then aligned against a ruminant chimera reference, as described in the screening section, using paired-end alignment parameters. Non-aligned reads were extracted, and duplicates were removed with PRINSEQ. Paired reads from multiple libraries belonging to the same sample were merged to create a single paired dataset per sample. Each merged dataset was then assembled with SPAdes using the --careful parameter to minimize mismatches and indels. Finally, the resulting contigs were mapped onto the modified Sheeppox virus reference genome using RAGTAG in scaffold mode (*126*).

The quality of the assembly was then assessed using CheckVv1.0.1 (*127*). Only samples classified as high quality according to CheckV and MIUViG criteria, with an estimated genome completeness greater than 90% and a minimum of 145 predicted genes, were retained for downstream analyses. Based on these criteria, seven samples (MYRZ180, RUB002, TCCL002, CL299, YG16, RF003, and ANC255) passed all quality filters (Table S12).

### Gene integrity assessment

To enable comparative analyses of gene integrity across the *Capripoxvirus* genus, particularly on the 9 genes reported to be inactivated in modern SPPV strains, we identified homologous sequences of the set of 156 LSDV reference genes in the 56 modern capripoxvirus genomes used in the Time-calibrated phylogeny of Capripoxviruses, recombination inference, and root estimation section and seven de novo-assembled ancient or historic genomes (MYRZ180, RUB002, CL299, TCCL002, YG16, RF003, and ANC255). Homologue detection was performed using a two-stage strategy combining Liftoff (*128*) and BLASTn (*129*). Gene annotations were primarily transferred from the LSDV reference genome to each target genome using Liftoff with minimum alignment fraction (-a) of 0.4 and minimum sequence identity (-s) of 0.4, allowing for annotation transfer across divergent sequences. Genes not identified by Liftoff were subsequently searched using BLASTn, where reference gene sequences were used as queries against target genome databases constructed using makeblastdb. BLASTn searches were performed with minimum percent identity ≥ 40% and minimum query coverage ≥ 30% of reference gene length.

Because gene inactivation in poxviruses can result from both single-nucleotide substitutions and frameshift mutations, accurate translation and reading frame assessment is critical. Each extracted gene sequence was translated using the standard genetic code, with ambiguous nucleotides (N) translated as X (unknown amino acid) to account for missing data in ancient DNA assemblies. The first canonical start codon (ATG) was searched within the first 60 nucleotides from the annotated start position. If no ATG was found in this region, the search was extended 30 nucleotides upstream. Translation was then performed following the reading frame of the identified start codon.

Stop codons were identified across translations, and sequences were classified into four categories: “Active”, if the sequence contained a start codon in the start region and no premature stop codons, with either a terminal stop codon within the last five amino acids or no stop codon detected; “Likely-active”, if the sequence met the criteria for active genes but contained ambiguous nucleotides (N) translated as X, accounting for assembly uncertainty in ancient DNA samples where some positions could not be confidently assembled but the overall gene structure appeared intact; “Inactivated”, if the sequence lacked a start codon or contained a premature stop codon more than five amino acids upstream of the sequence end; and “Not found”, for loci that were not successfully annotated in the target genome through either Liftoff or BLASTn rescue. For ancient sequences, gene inactivation status was manually verified by inspection using Geneious (https://www.geneious.com).

### Parallel or ancestral gene inactivation analysis

To evaluate whether the parallel inactivation of 8–9 genes in the GTPV and SPPV lineages could result from a single ancestral inactivation event, we searched for shared inactivating mutations between the two lineages. If gene inactivation occurred in their common ancestor, at least one identical inactivating mutation should be detectable in both GTPV and SPPV. The nine inactivated genes were aligned in-frame using MACSE v2.07 (*130*), with the LSDV reference sequence used as the functional reference to preserve the ancestral coding state. MACSE is a codon-aware aligner that accounts for frameshifts and premature stop codons, which are the predominant mechanisms of gene inactivation in poxviruses. Frameshift mutations and premature stop codons (excluding the terminal stop codon) were considered inactivating events. Each mutation was annotated according to its type and position within the predicted protein sequence. The resulting set of inactivating mutations was then compared across all GTPV and SPPV strains to identify mutations shared between lineages, which would support an ancestral origin or parallel origin of gene inactivation.

### Positive selection

To test for evidence of positive selection in our dataset between our ancient genomes and modern we used the codeml program implemented in PAML (*131*). The exact same methods as described above have been used to find homologous genes in the same SPPV modern datasets used for the Maximum Likelihood building and the de-novo reconstructed ancient genomes but by using SPPV reference (NC_004002.1) genes instead. Then genes have been then aligned as described above with MACSE and cleaned and placed in frame by using the codon-msa modules for Hyphy (https://github.com/veg/hyphy-analyses/tree/master/codon-msa). Following data preparation, the branch-site test in PAML was used (model=2; NSsites=2; fix_omega=0/1; omega = 1), by annotating modern sequences and ANC255 (1917 CE) as foreground lineages and the rest of the ancient sequences as background lineage. Pairs of log-likelihood values for each locus were compared using a likelihood ratio test (2ΔlnL) with one degree of freedom (d.f. = 1). Final P-values were obtained by dividing the original P-values by two, following the procedure described in the PAML manual. Multiple testing correction was performed using the Benjamini–Hochberg method. No tests were positive after multiple testing corrections; we report these negative results in Table S6.

## Acknowledgements

We acknowledge the following persons and institutions without which this work and manuscript would not be possible: Ciarán O’Connor, Deborah Diquelou, Victoria Mullin, Valeria Mattiangeli, Beinecke Rare Book and Manuscript Library (Speculum historiale samples), The Master and Fellows of Trinity College, Cambridge (TCCL02-10), Parker Library, Corpus Christi College, Cambridge (CL299), The Syndics of Cambridge University Library, Philippa Hoskin, Nicolas Bell, Suzanne Paul, St. Florian Abbey (Austria), Mag. Harald Ehrl Can. Reg., Project MALADI, chaire SHS Île-de-France, François Gentili (INRAP), Ariane Düx, Lilli Gralla, Annika Graaf-Rau, Pip Willcox, Lambeth Palace Library, the Borthwick Institute for Archives, Christopher C Webb, Catherine Firth, Alison Fairburn, the Austrian National Library, Christa Hofmann.

## Author contributions

Conceptualization: KGD

Supervision: KGD, SC

Investigation: LH, LS, RG, AS, LR, JL, LA, EL, MB, SF, MDT, JV, JS, KGD

Formal analysis: LH, KGD, TD

Data curation: LH, MDT, JS

Resources: BR, HW, RH, PE, MTJW, KR, EN, MMA, MR, LCV, DB, JS, MC, DVP, AB

Writing—original draft: KGD, LL

Writing—review & editing: KGD, LL, EL, JS, MC, AB, FK, CM, RH, DB, all authors

## Data and materials availability

Reads aligned to sheeppox virus and host mitochondrial (and secondary contaminant) genomes are available as non-filtered BAM files at ENA under accession PRJEB107108. Codes used in this study and BEAST input files are available at https://github.com/LouisLhote/Sheeppox_aDNA. Previously published modern capripoxvirus datasets are listed in Table S6 together with the associated accession numbers and available metadata. Any additional information required to reanalyze the data reported in this paper is available from the lead contact upon request. All data are available in the main text or the supplementary materials.

