## Supplementary figures for "3,500 years of sheeppox virus evolution inferred from archaeological and codicological genomes"

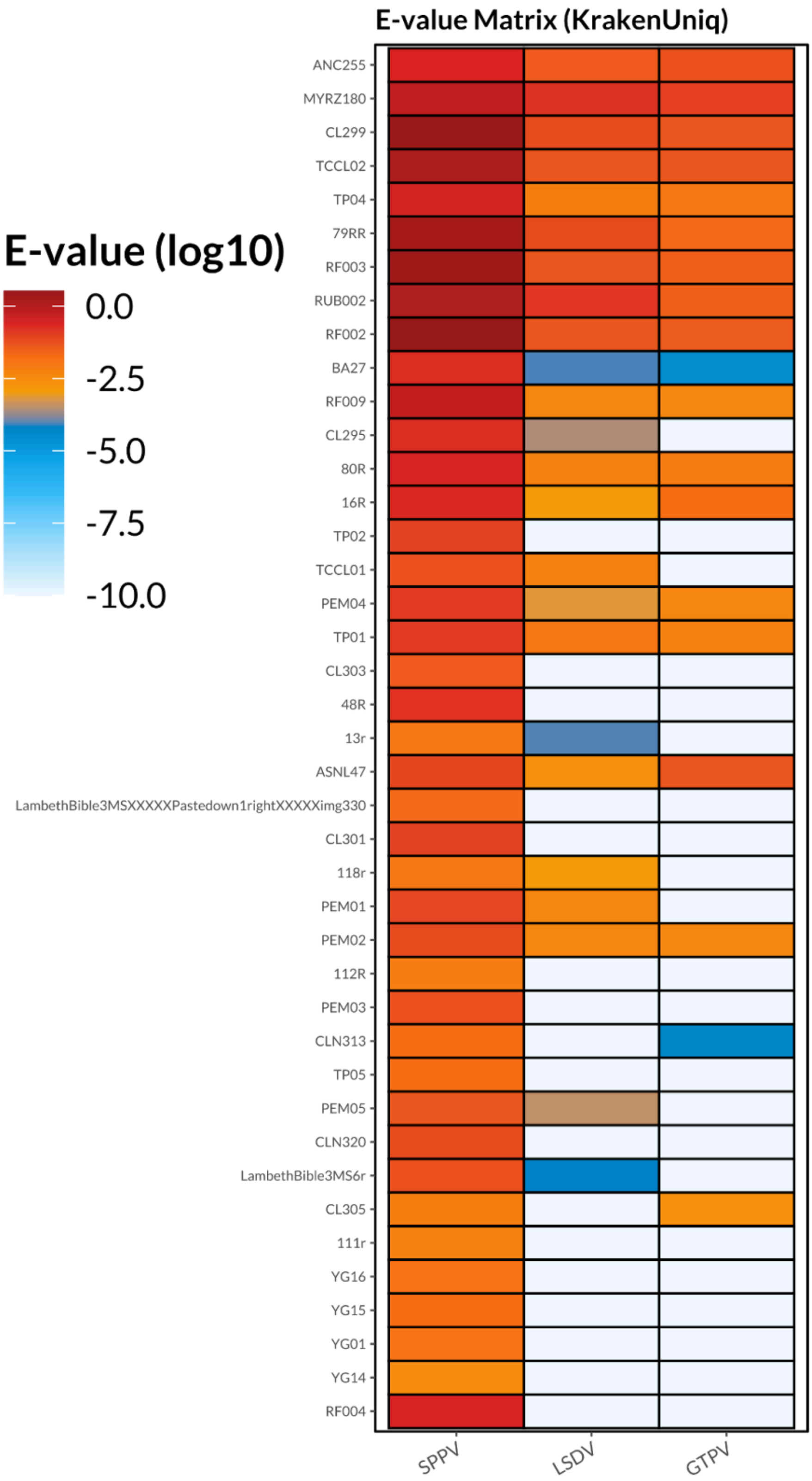


**Supplementary Figure 1.** *Krakenuniq E-value for Capripoxviruses.* Heatmap showing log10-transformed E-values for Capripoxviruses-positive samples and viral species (columns). Warmer colors indicate high E-values, whereas cooler colors indicate low E-value.

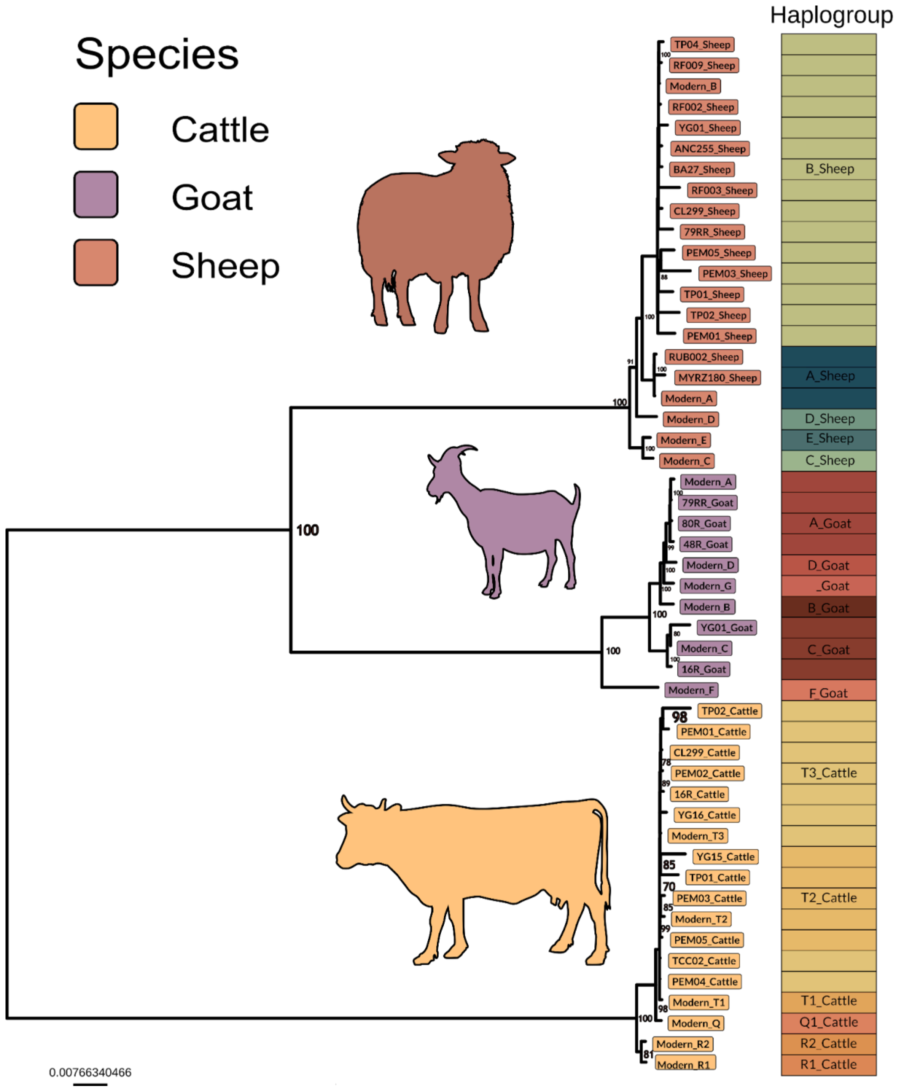


**Supplementary Figure 2**. *Host-associated Mitochondrial DNA Maximum Likelihood Phylogeny.* Maximum likelihood mitochondrial tree including ancient and modern haplogroup references, colored by species (sheep, goat, cattle). Bootstrap support values are shown for nodes with bootstrap values > 70. Major species clades are clearly separated, and samples are further annotated by mitochondrial haplogroups (right panel). Silhouettes illustrate species groupings.

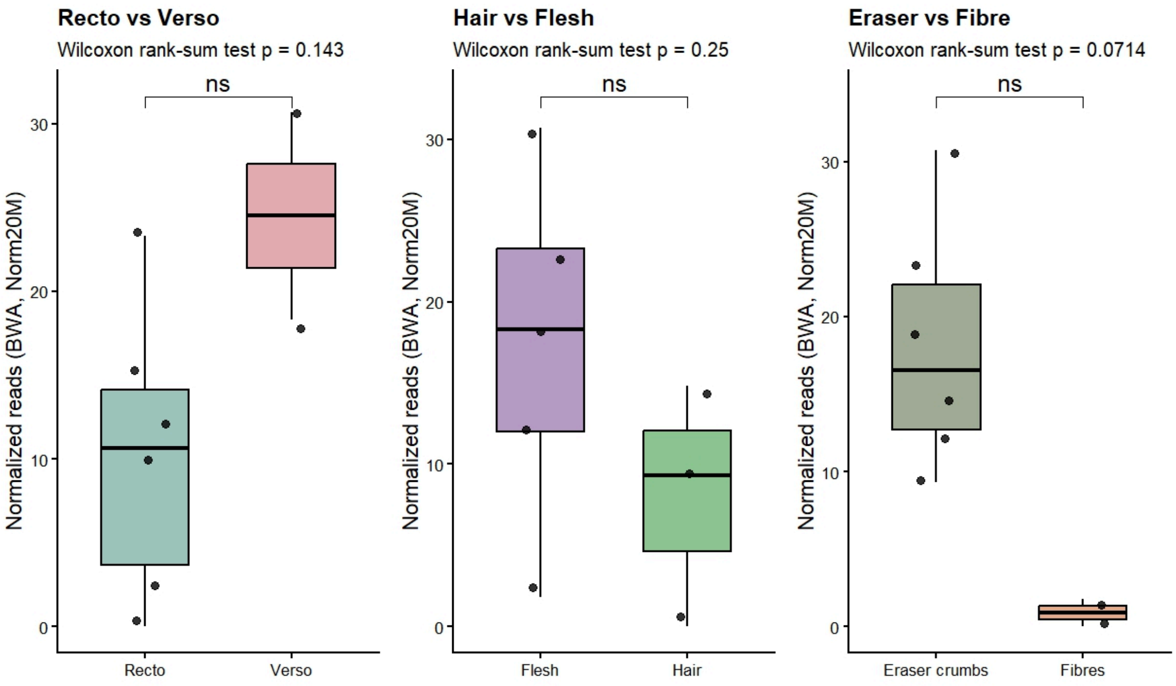


**Supplementary Figure 3.** *Comparison of normalized SPPV reads from localized sampling of sample TCCL2.* Boxplox showing SPPV reads aligned to the sheeppox reference genome and normalized to 20 million reads for three comparisons: recto vs verso, hair vs flesh, and eraser crumbs vs fibres. Dots represent individual samples. Statistical differences between groups were computed using an unpaired Wilcoxon rank-sum test. P-values are reported above each plot (p *< 0.05; p** < 0.01; ***p < 0.001; ns, not significant)

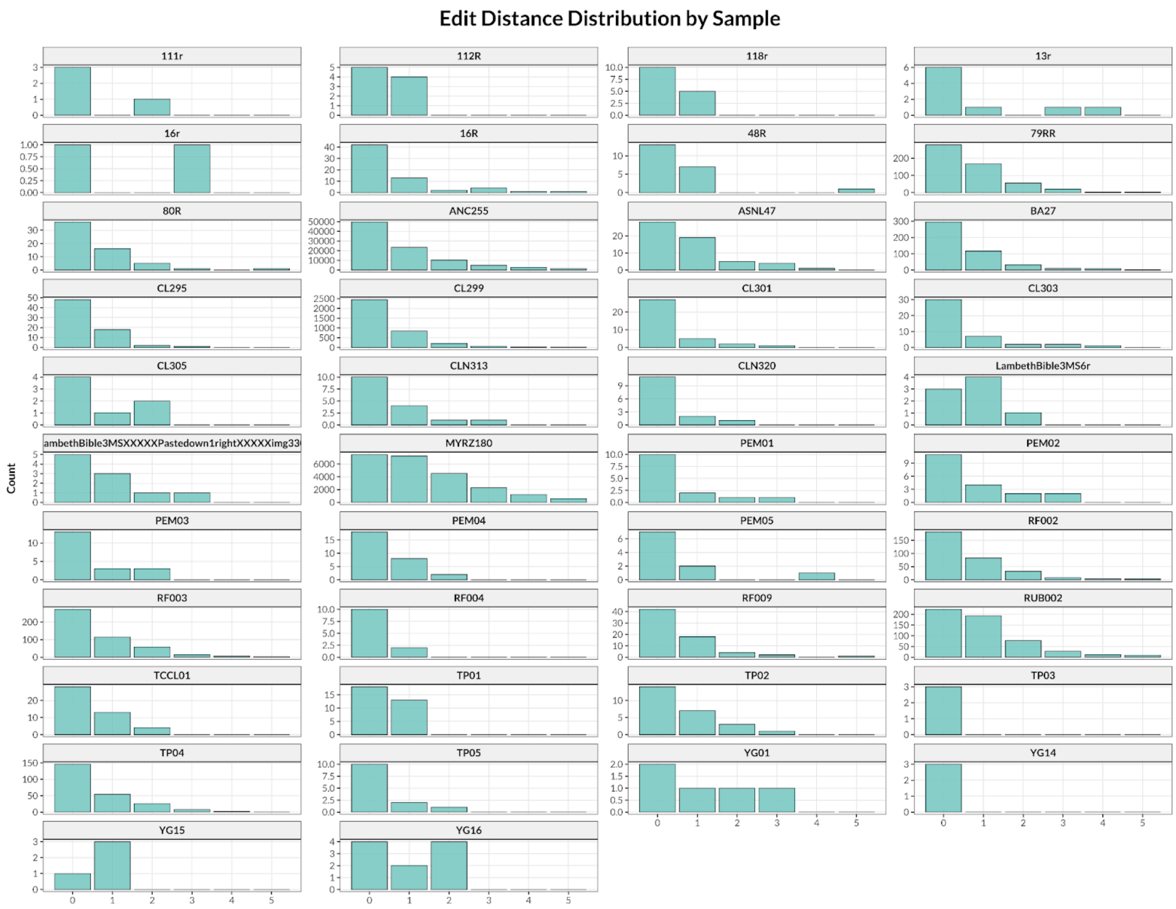


**Supplementary Figure 4**. *Edit distance distribution of Capripoxvirus-positive samples relative to the SPPV reference genome.* For each screened Capripoxvirus-positive sample, the distribution of edit distances to the SPPV reference genome is shown. The first five edit distance values are displayed for each sample.

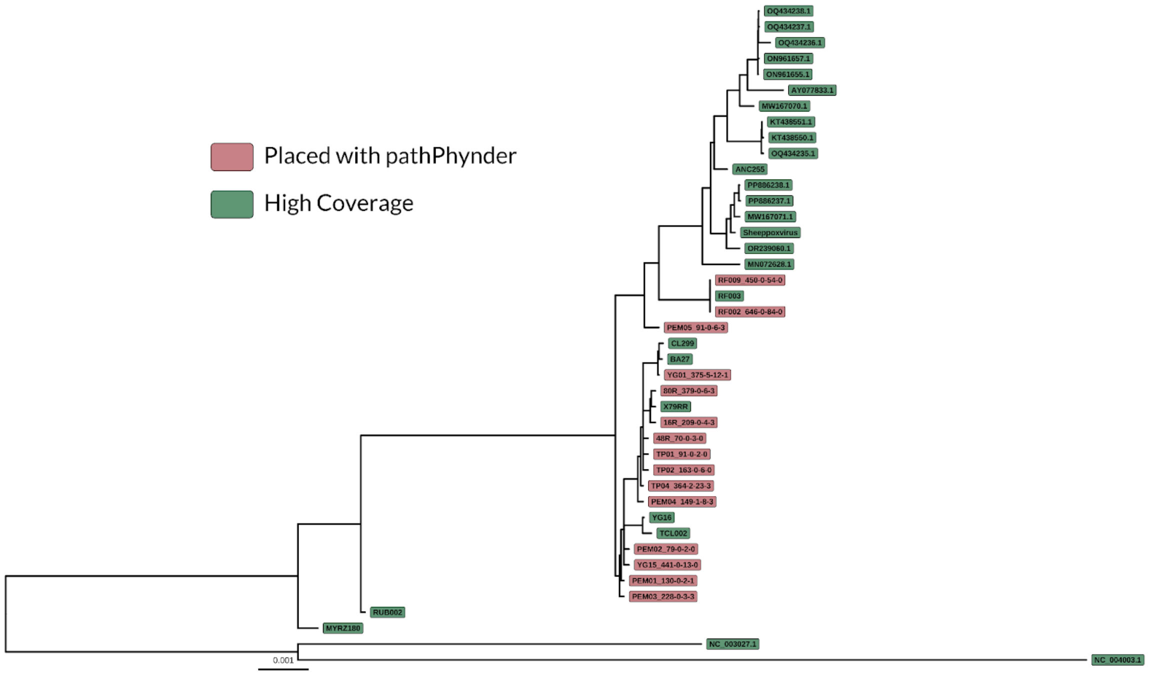


**Supplementary Figure 5.** *SPPV Maximum likelihood phylogeny with low-coverage samples placed using pathPhynder.* The maximum likelihood tree is the same as that shown in Figures 2A and 2B. Low-coverage samples placed using pathPhynder are highlighted in red, whereas high-coverage samples are shown in green. For each low-coverage sample, the corresponding pathPhynder SNP-path placement details are displayed. Further information on the SNP-path placement methodology is available at https://github.com/ruidlpm/pathPhynder.

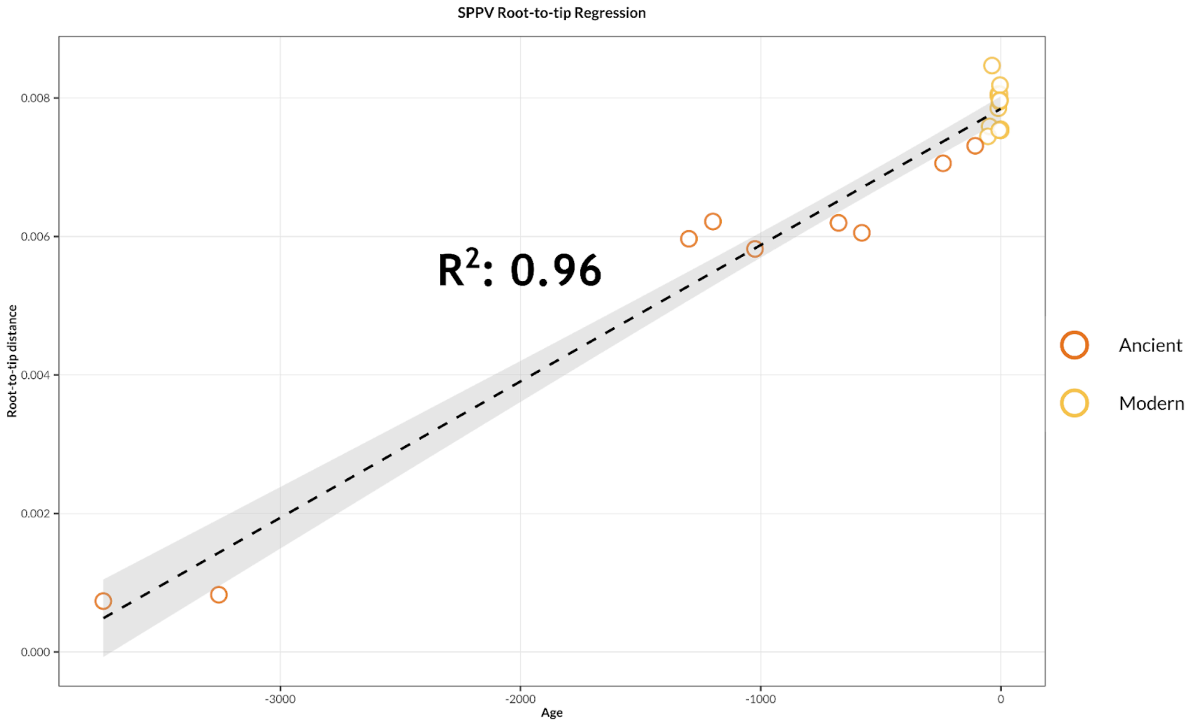


**Supplementary Figure 6.** *Root-to-tip regression of SPPV samples as function of age.* Root-to-tip genetic distances are plotted against sample age, expressed as time before present. The dashed line represents the linear regression, and the $R^2$ is shown.

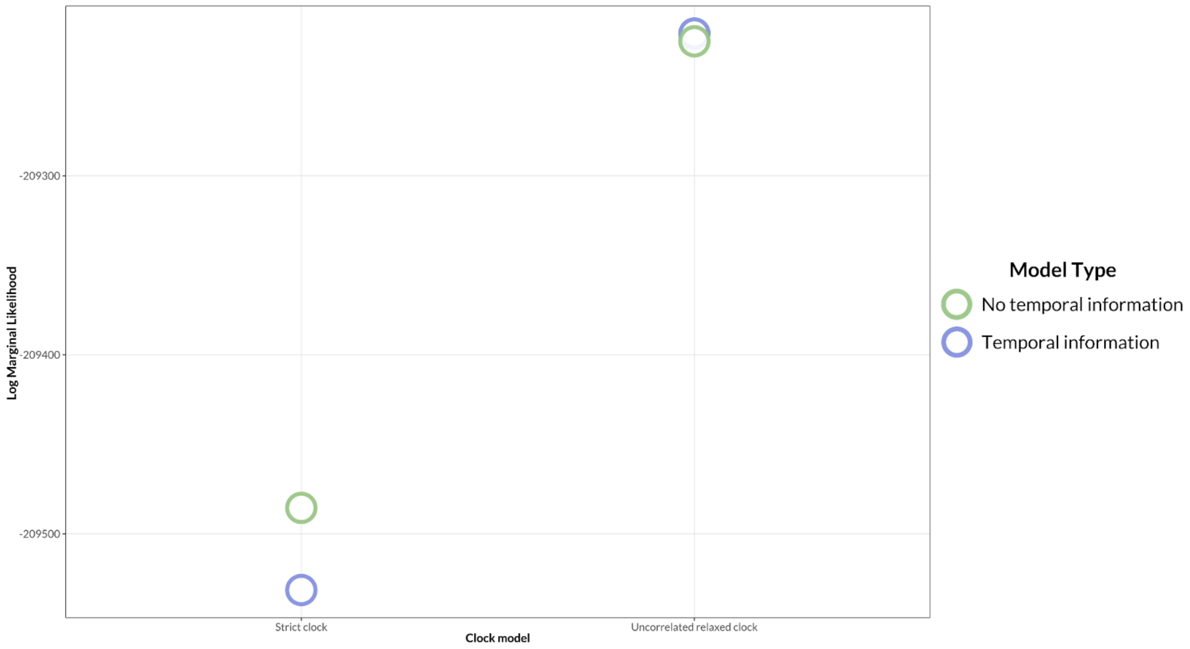


**Supplementary Figure 7.** *Log marginal likelihoods of SPPV BEAST models.* The log marginal likelihood of each of the four BEAST models tested is shown as a single point. Models incorporating sample temporal information are shown in blue, whereas models without temporal signal are shown in green. Models are arranged along the x-axis according to the molecular clock model used.

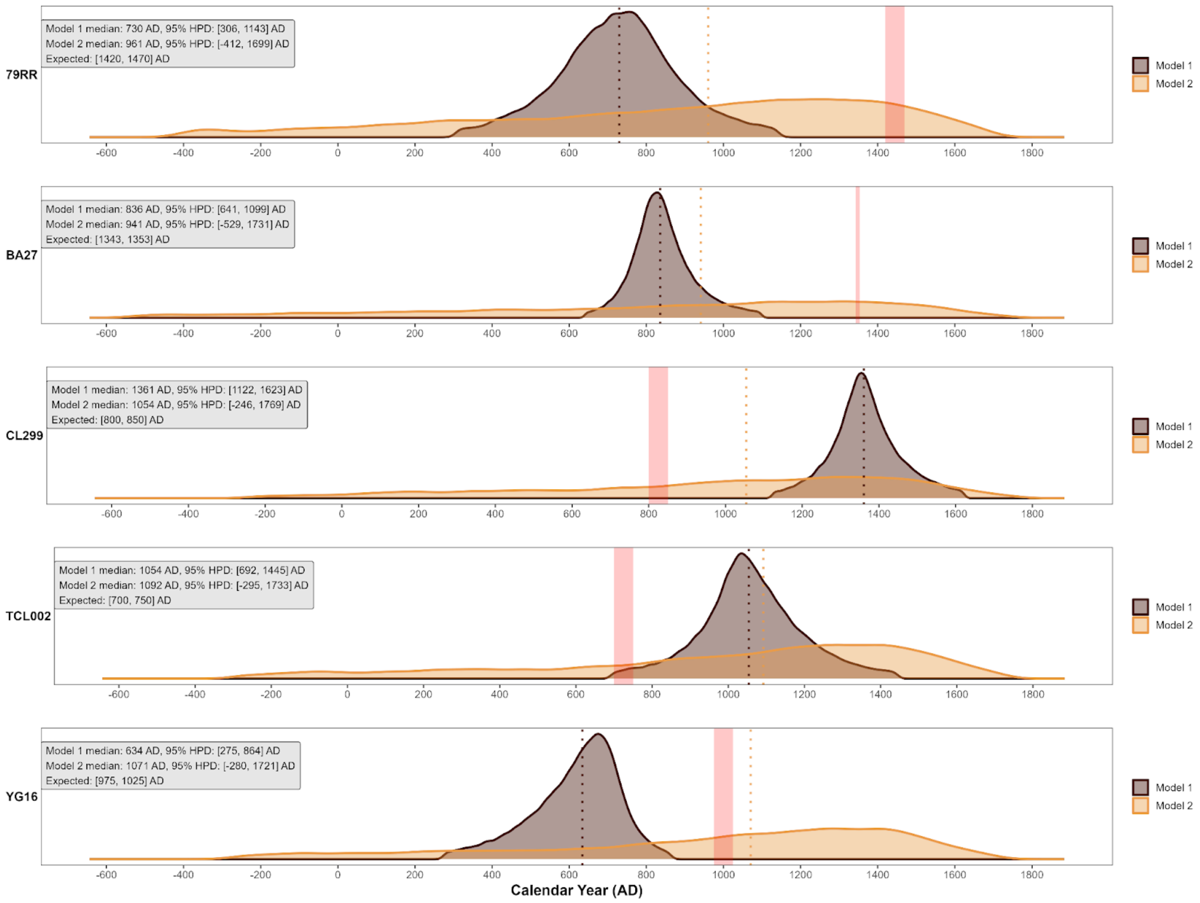


**Supplementary Figure 8.** *Tentative tip dating of SPPV manuscript-derived samples.* For each parchment-derived sample, two models of tip dating are shown: Model 1, in which one of the five parchment-derived sample ages is replaced by a uniform prior spanning the temporal range of the tree; and Model 2, in which all parchment-derived samples are excluded from the analysis except for one assigned a uniform prior spanning the temporal range of the tree. For each parchment-derived sample, the 95% highest posterior density age intervals are shown in grey (Model 1) and orange (Model 2), with a red line representing the historically inferred age shown for comparison. The median of each 95% HPD interval is represented by a dashed line, and median values are reported in the grey box in the top-left corner of each plot. The log marginal likelihood of each of the four BEAST models tested is shown as a single point. Models incorporating sample temporal information are shown in blue, whereas models without temporal signal are shown in green. Models are arranged along the x-axis according to the molecular clock model used.

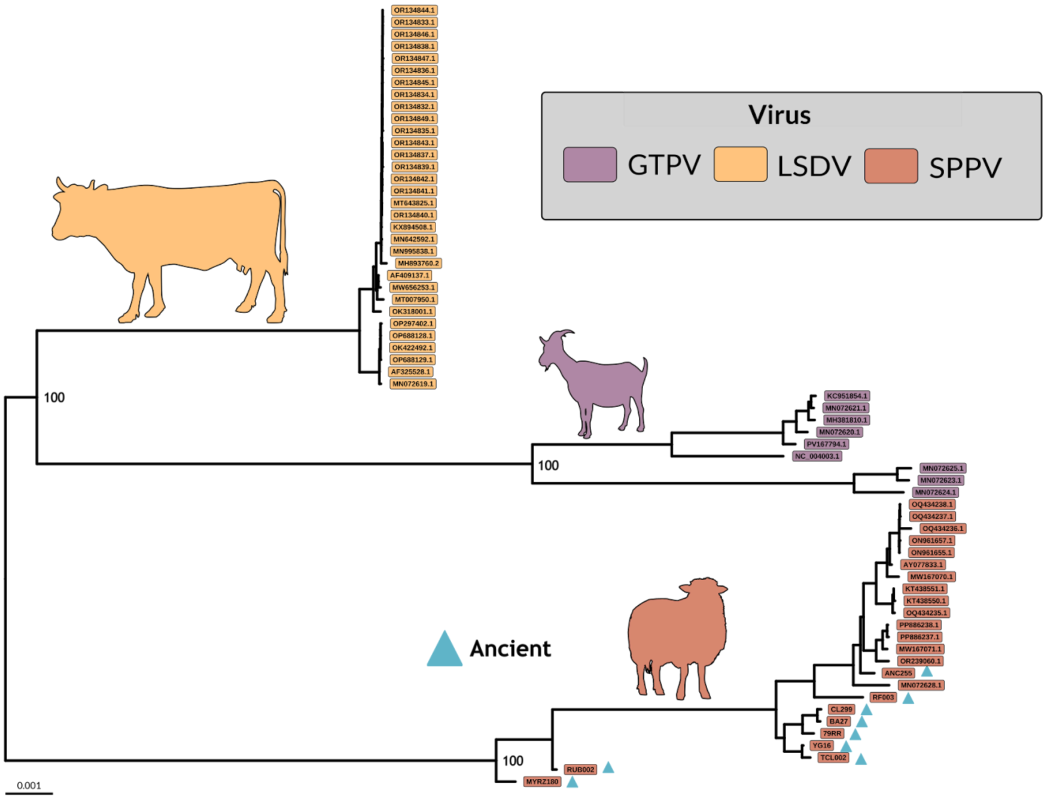


**Supplementary Figure 9.** *Maximum likelihood tree of Capripoxvirus dataset*. Each sample is colored according to its viral species, and silhouettes of the corresponding typical hosts are shown in matching colors. Ancient samples are highlighted with blue triangles. Bootstrap support values for nodes defining each species are indicated at the corresponding nodes.

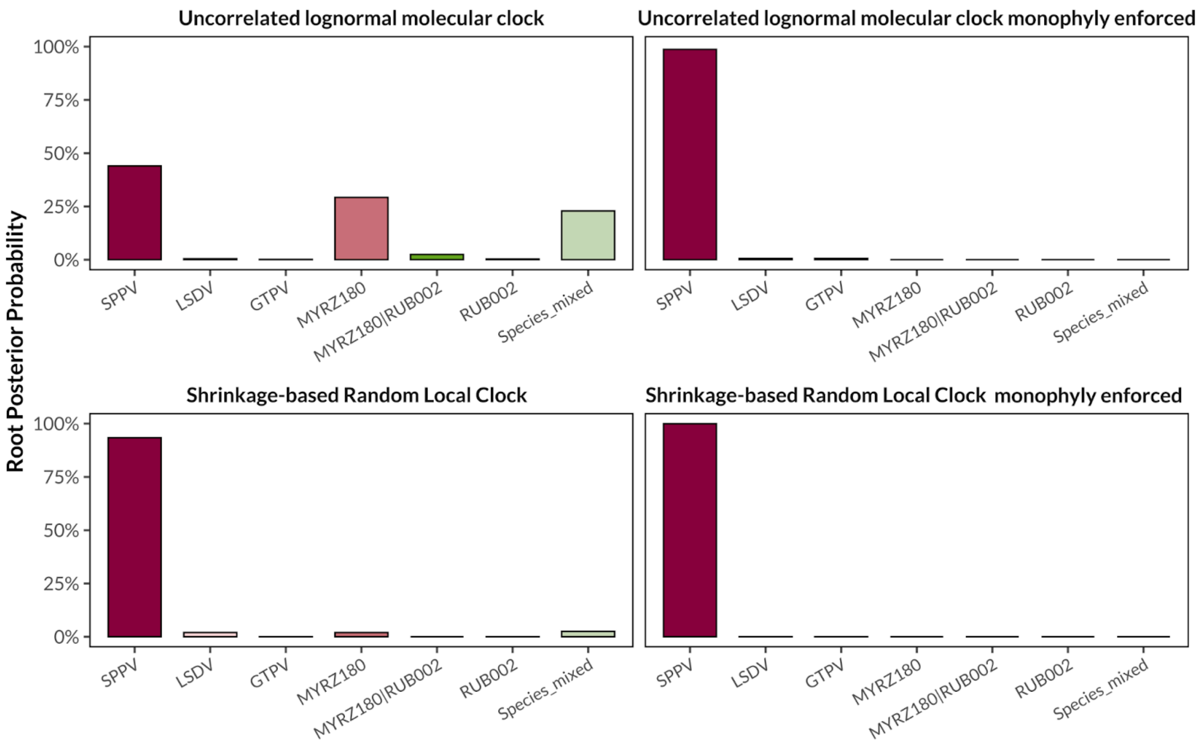


**Supplementary Figure 10.** *Root posterior probabilities of Capripoxvirus BEAST models with and without monophyly constraints.* Posterior probabilities for each possible root placement are shown for each molecular clock model, analyzed with and without enforcement of species monophyly. “Species mixed” denotes cases in which BEAST inferred a root composed of samples from different capripoxvirus species. Colors indicate corresponding root positions and are consistent across models.

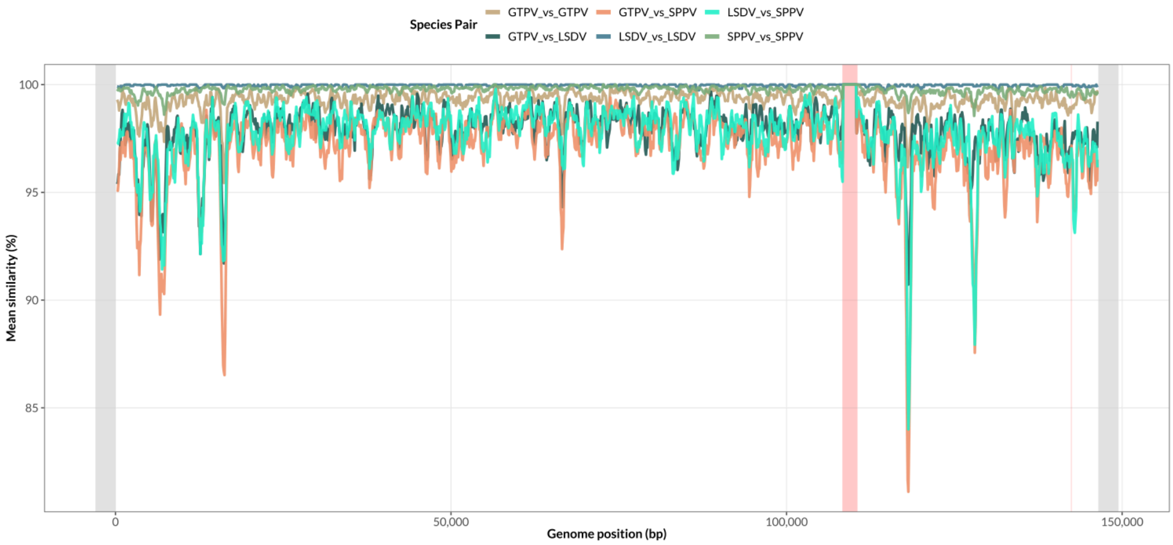


**Supplementary Figure 11.** *Sliding-window pairwise similarities along the Capripoxvirus genome alignment.* Mean inter- and intra-species pairwise similarity values computed in sliding windows (500 bp, 100 bp steps) along the Capripoxvirus genome alignment are shown in different colors according to the species pair. Genomic regions excluded from the analysis at the alignment edges are highlighted in light grey, and putative recombinant regions are highlighted in red.

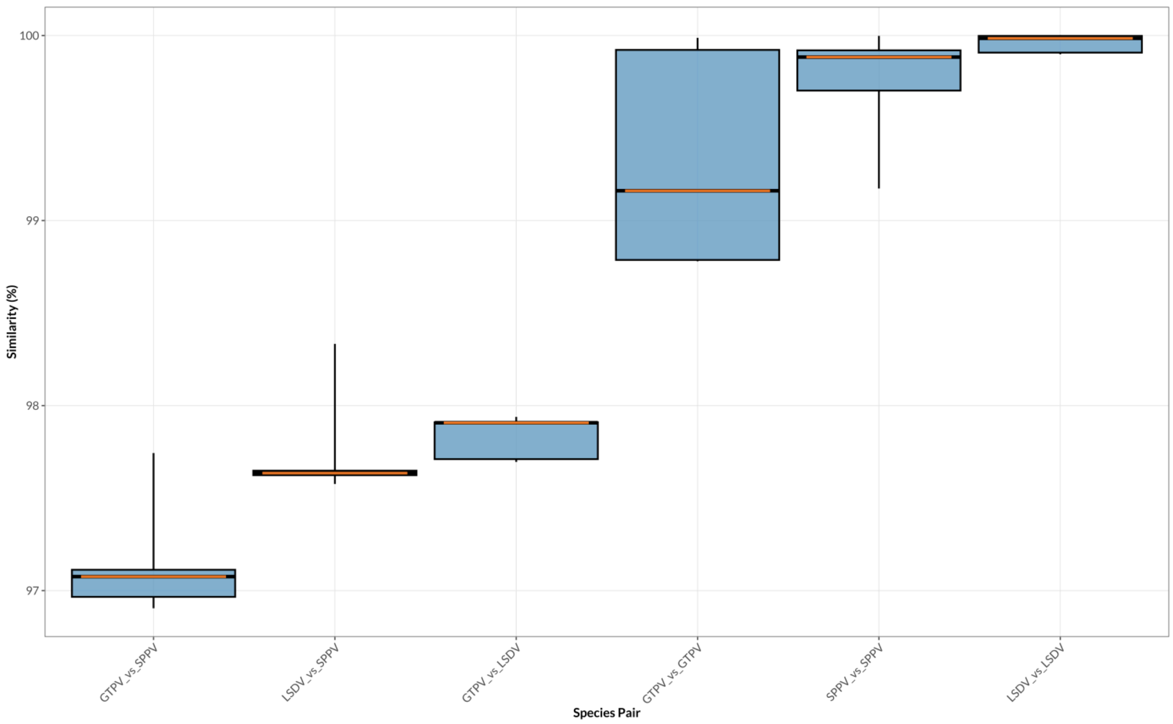


**Supplementary Figure 12.** *Genome-wide pairwise similarities among Capripoxvirus clades.* The distribution of genome-wide pairwise similarity values between individual samples from each Capripoxvirus clade pair is shown in blue. The median similarity value for each distribution is indicated in orange.

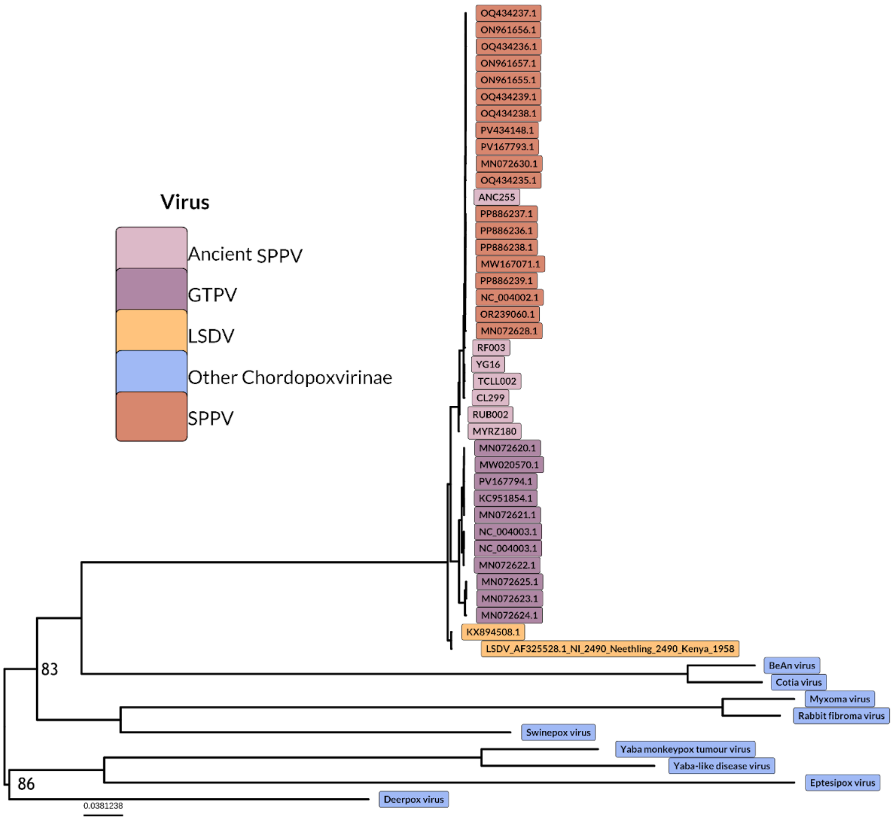


**Supplementary Figure 13.** A maximum likelihood phylogeny of the SPPV samples with GTPV, LSDV, and the nine most closely related chordopox species based on the KX894508.1 core genome coordinates corresponding to CDSs LSDV027-LSDV123. The 30 modern SPPV genomes are shown by red tips, 15 GTPV by blue tips, two representative LSDV by green tips, and the seven ancient SPPV and nine chordopox species by black tips. The data for the nine chordopox species came from: deerpox virus NC_006966.1, Yaba-like disease virus NC_002642.1, Yaba monkeypox tumour virus NC_005179.1, Cotia virus NC_016924.1, BeAn virus NC_032111.1, Rabbit fibroma virus NC_001266.1, Myxoma virus NC_001132.2, Swinepox virus NC_003389.1, and Eptesipox virus NC_035460.1. The tree is midpoint rooted and only Bootstrap support values below 90% are displayed.. The scale bar shows substitutions per site. Adding in the other 60 published LSDV WT core genomes had no effect on the topology (data not shown), and so two are shown here for visual clarity.

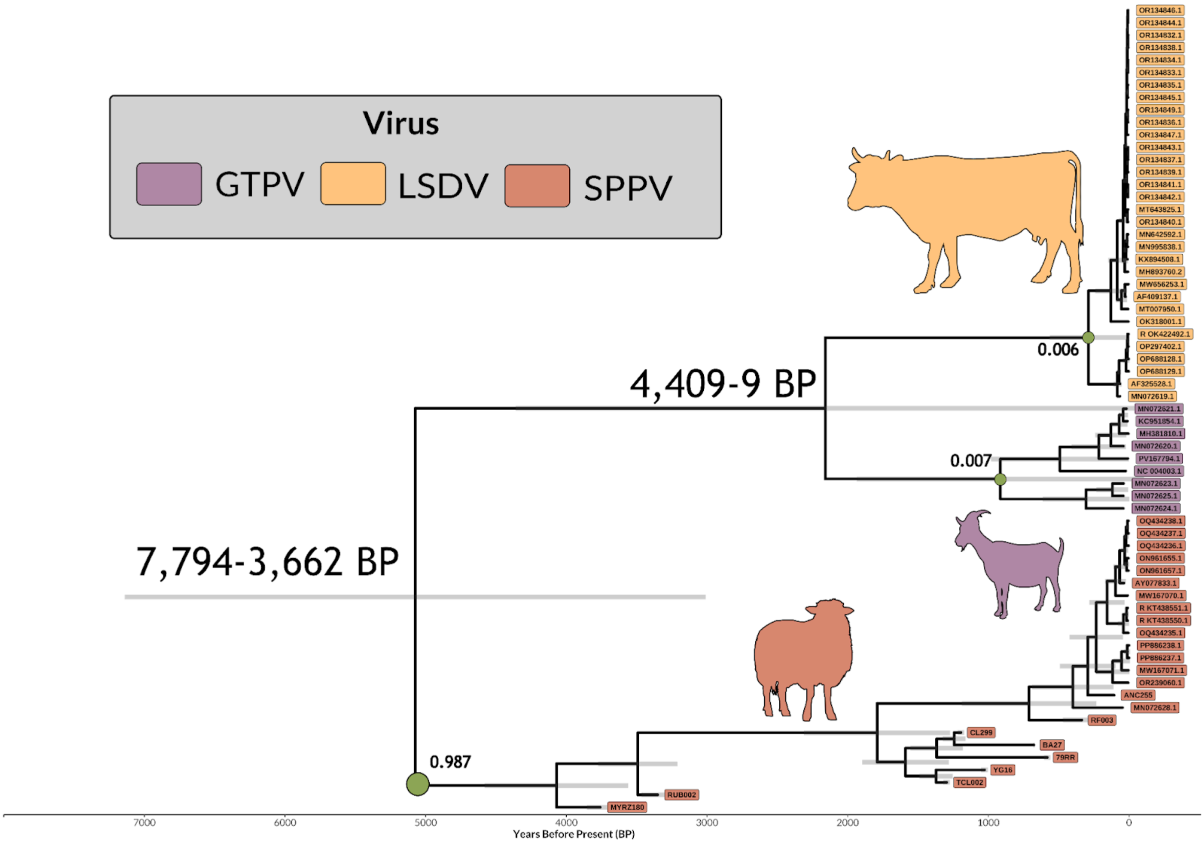


**Supplementary Figure 14.** *BEAST phylogeny of the Capripoxvirus dataset inferred under an uncorrelated lognormal relaxed molecular clock model.* Each sample is colored according to its viral species, and silhouettes of the corresponding typical hosts are shown in matching colors. The time scale is shown in years before present.

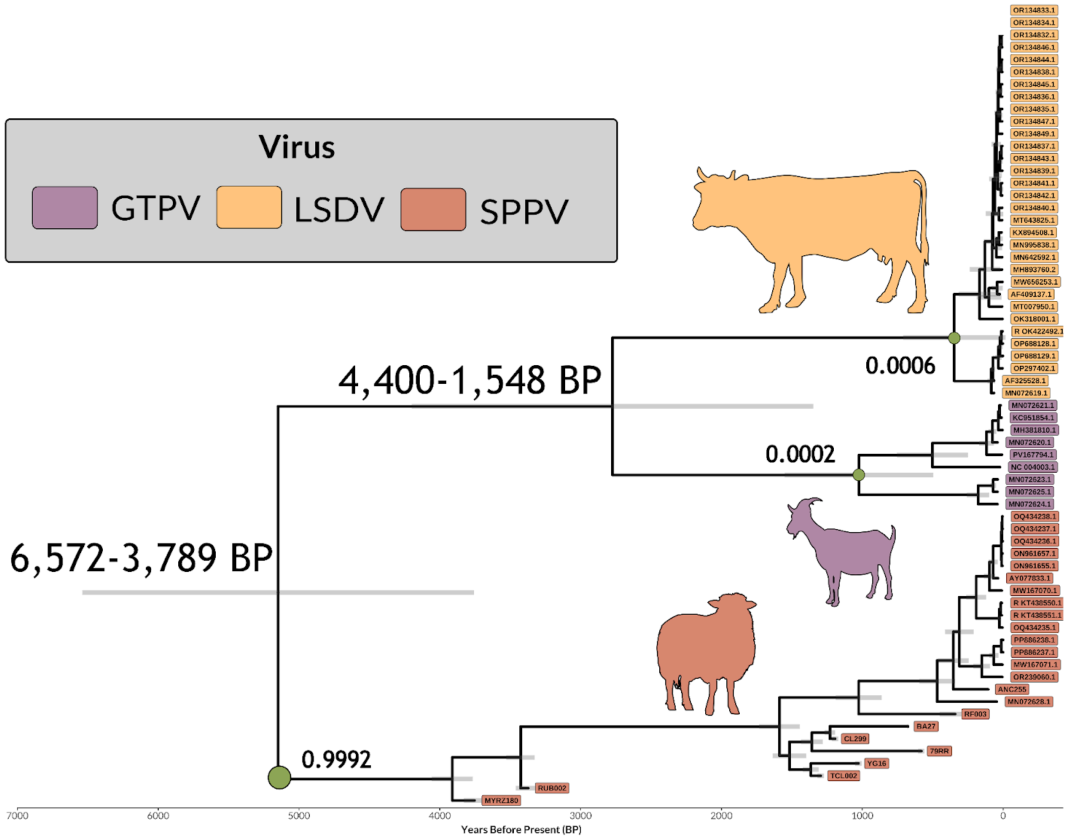


**Supplementary Figure 15:** *BEAST phylogeny of the Capripoxvirus dataset inferred under an Shrinkage-based Random Local Clock model.* Each sample is colored according to its viral species, and silhouettes of the corresponding typical hosts are shown in matching colors. The time scale is shown in years before present.

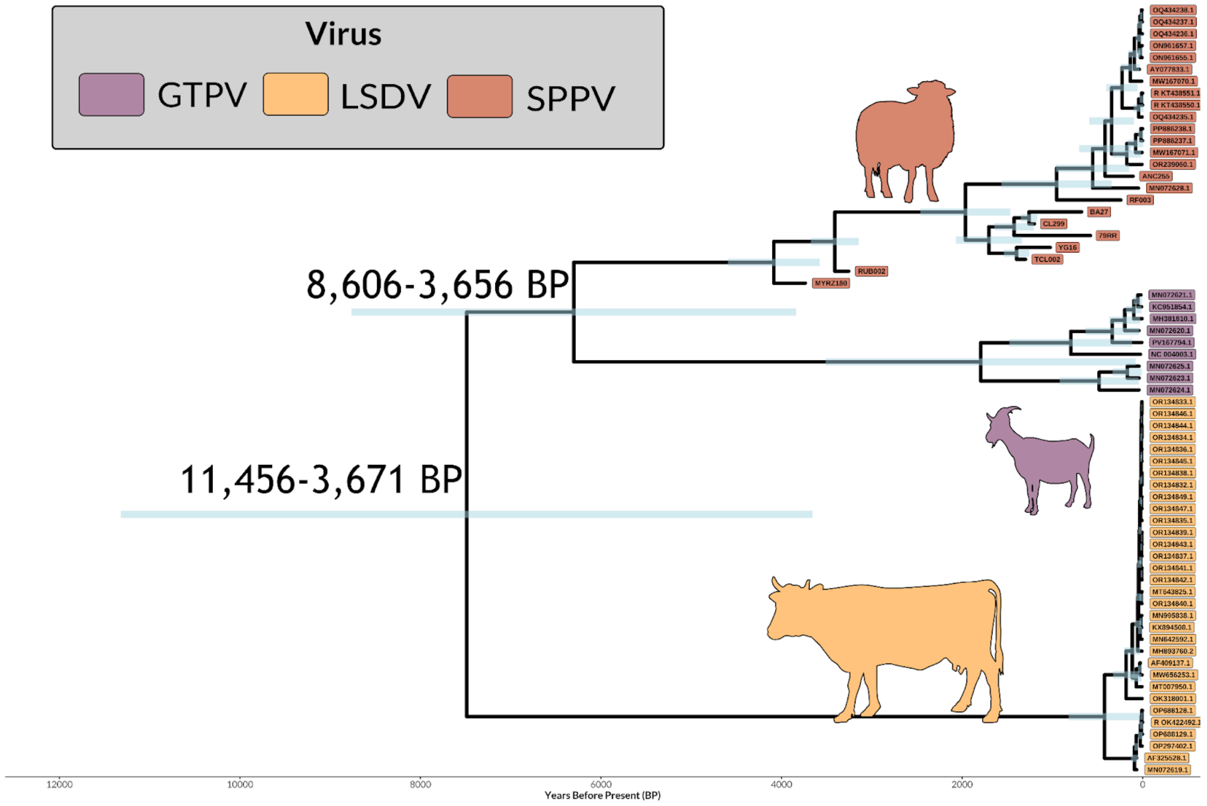


**Supplementary Figure 16.** *BEAST phylogeny of the Capripoxvirus dataset inferred under an uncorrelated lognormal relaxed molecular clock model with LSDV rooting enforced.* Each sample is colored according to its viral species, and silhouettes of the corresponding typical hosts are shown in matching colors. The time scale is shown in years before present.

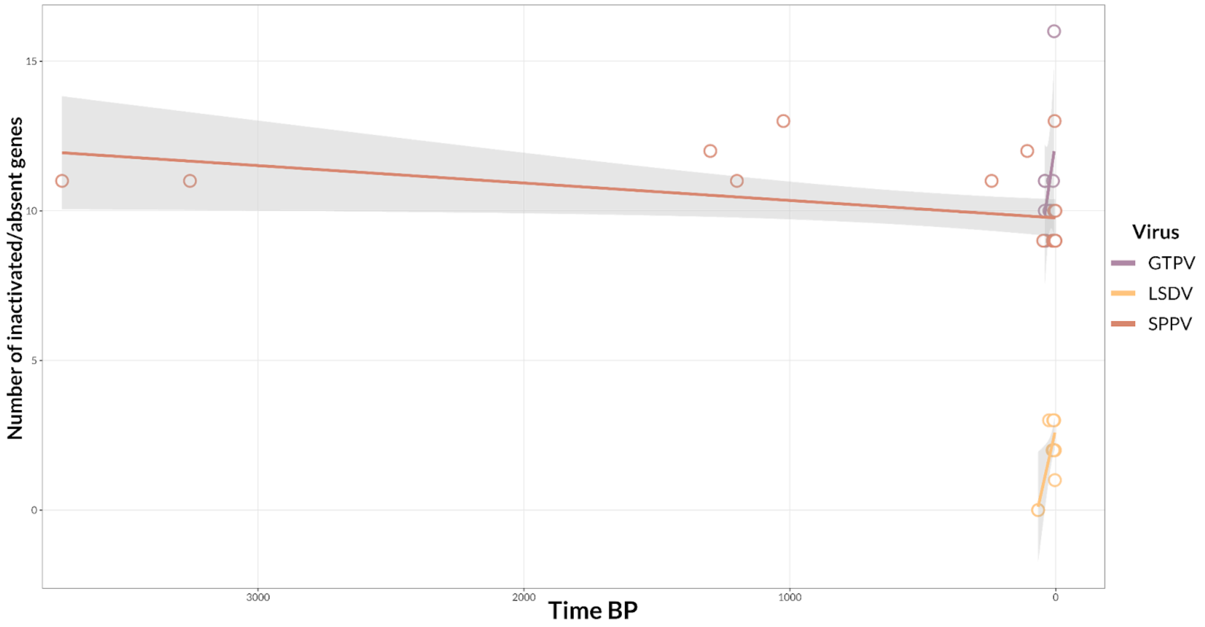


**Supplementary Figure 17.** *Capripoxvirus gene inactivation through time.* Each capripoxvirus sample is represented by a point, with sample age (years before present, BP) shown on the x-axis and the number of gene inactivations in the genome on the y-axis. Points are colored according to viral species, and a regression line is fitted for each viral group.

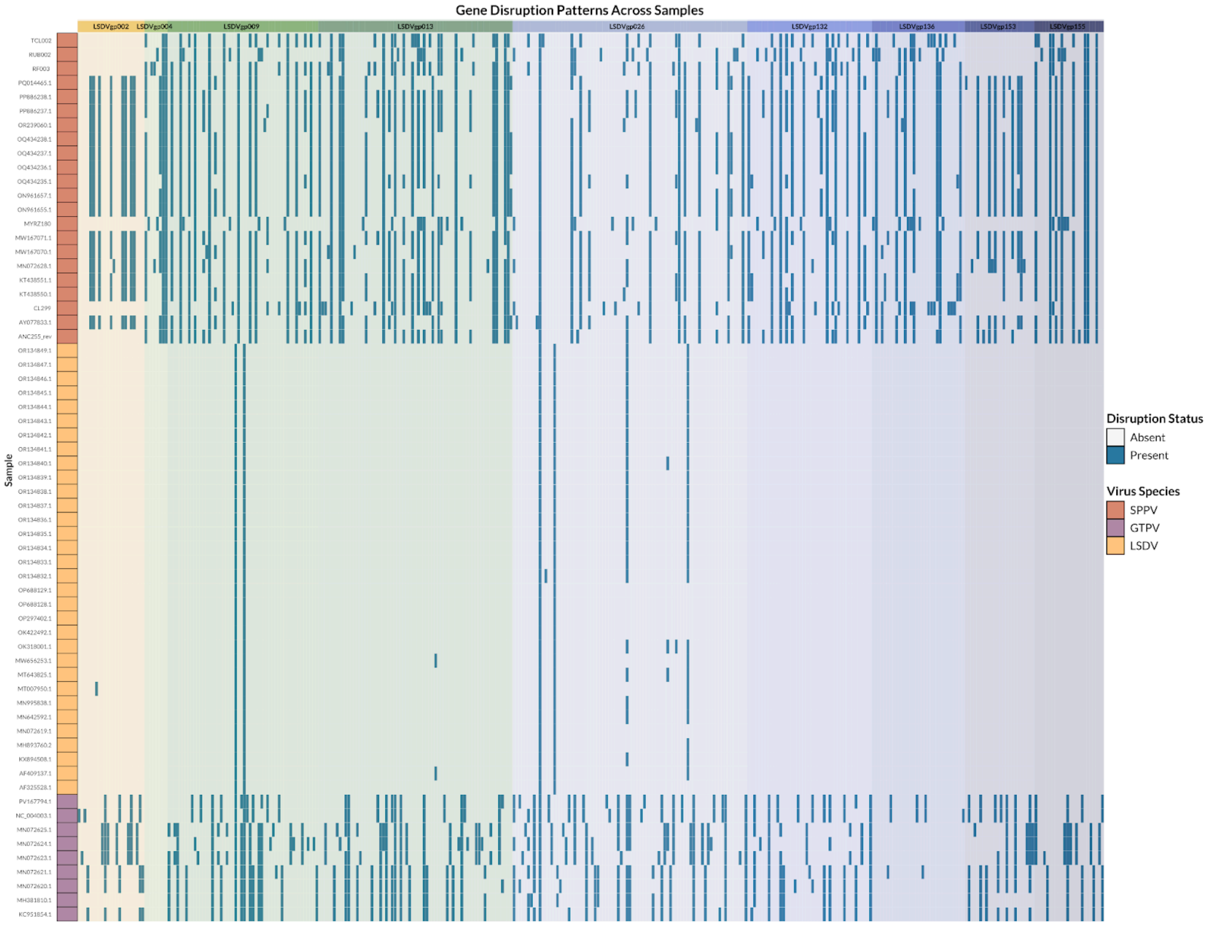


**Supplementary Figure 18.** *Gene disruptions in inactivated Capripoxvirus genes.* The heatmap shows the presence and absence of gene disruptions in inactivated capripoxvirus genes across all samples. Genes are arranged by background color, and viral samples are shown in rows and colored according to species. Disruptions are indicated in blue, whereas absence of disruption is shown in white. Details in Supplementary Table 11.

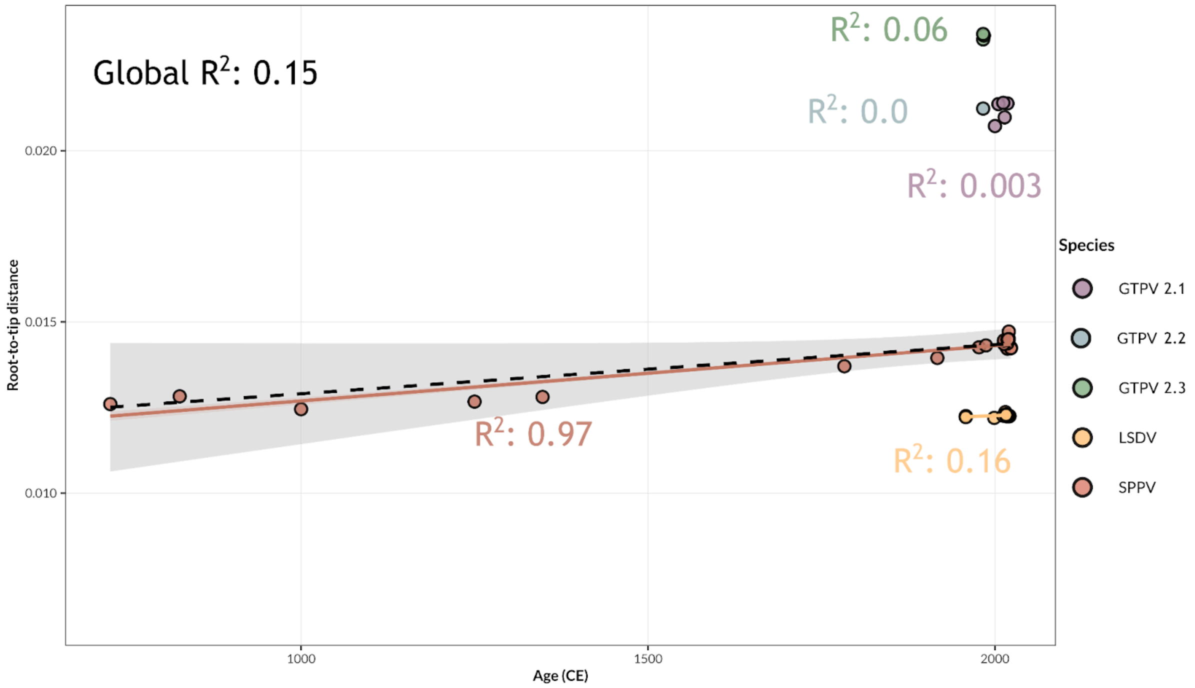


**Supplementary Figure 19.** *Root-to-tip regression of Capripoxviruses samples as function of age.* Root-to-tip genetic distances are plotted against sample age, expressed as time before present. The dashed line represents the Capripoxvirus global linear regression, and the $R^2$ is shown. Samples are colored by viral clade origin and species specific $R^2$ are shown.

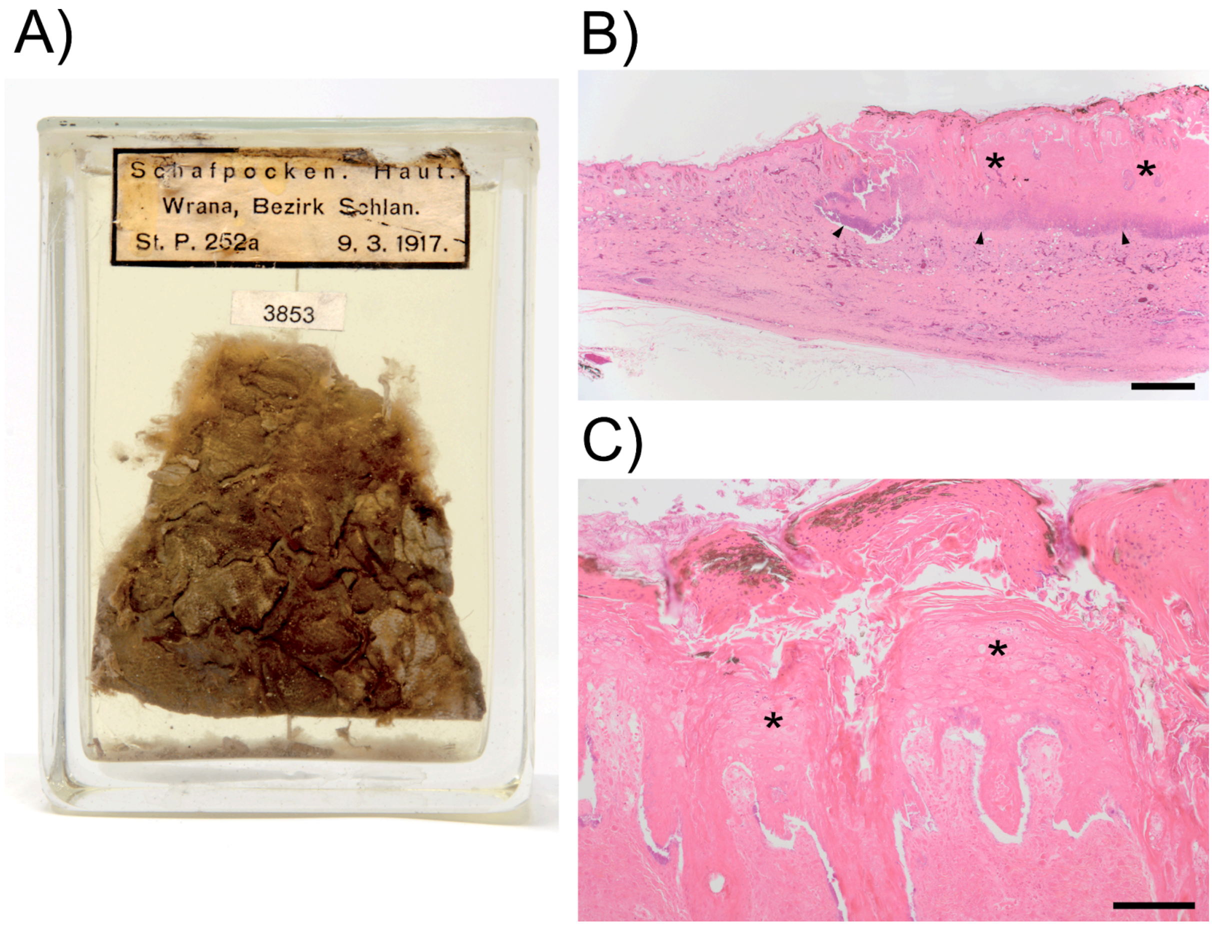


**Supplementary Figure 20.** *Gross and histological description of sample ANC255.* A) Archived skin specimen from a sheep infected with sheeppox virus. Typical lesions are irregular confluent papules and nodules leading to thickening of the skin and clear demarcation towards unaffected areas of the sparsely haired skin. B) Histological section of the skin specimen seen in A. The thickened areas of the skin result from deep necrosis of the dermis and hyperplastic epidermis (asterisks). A leucocytic demarcation line is seen in the superficial subcutis (arrowheads). Haematoxylin-Eosin, bar = 1 mm. C) Closer view of the necrotic area of B. Ballooning degeneration is seen in the epidermal epithelium (asterisks) and the surface layers show severe parakeratotic hyperkeratosis. Haematoxylin-Eosin, bar = 160 µm.
